# Expanding the therapeutic window of bacteria for cancer therapy

**DOI:** 10.64898/2026.10.27.761579

**Authors:** Jongwon Im, Meghna Komaranchath, Sophia Windemuth, Hannah Kim, Xiaoyu Hu, Arjun Adapa, Myra Ashraf, Katherine A. Cummins, Courtney Coker, YoungUk Jang, Hayoung Jin, Kam W. Leong, Ronny Drapkin, Jeffrey N. Bruce, Peter Canoll, Tal Danino

**Author notes:** These authors contributed equally to this work (Jongwon Im, Meghna Komaranchath).

## Abstract

Engineered bacteria enable self-amplifying therapeutic delivery and immune activation, yet these same features drive systemic toxicity and off-target growth. Here, we create a “tumorobe” that overcomes these limitations by integrating reduced immunogenicity, tumour-restricted growth, and enhanced payload release. Specifically, we identify lipopolysaccharide modifications that reduce immunogenicity while preserving fitness, restrict growth to tumours using a hypoxia–lactate “AND” gate, and enhance extracellular payload release via lipoprotein deletion. This strain expands the maximum tolerated dose in a route-dependent manner by up to 1,000-fold. Across multiple murine models, including colorectal cancer and glioblastoma, the strain drives robust antitumour efficacy as both a standalone therapeutic and an immunotherapeutic delivery platform, achieving durable complete responses in up to 75% of colorectal tumours when armed with a cytokine payload. Together, these results establish a broadly applicable platform for safe and effective bacterial cancer therapy.

---

Tumours create ecological niches that are inhospitable to normal host cells yet permissive to microbial life. Within these hypoxic and immunosuppressed environments, diverse obligate and facultative anaerobic species have been identified across human cancers and linked to modulation of therapeutic response^1–3^. These observations motivate the development of microbial therapies, but naturally occurring species rarely integrate core design requirements such as tumour selectivity, low immunogenicity, and genetic tractability within a single organism.

As a result, efforts have focused on genetically modifying model organisms to achieve ideal characteristics, most often rapidly-growing Gram-negative facultative anaerobes such as *Salmonella* and *Escherichia coli*, although engineered commensals from other niches have also demonstrated antitumour potential^4–6^. Yet, their intrinsic activation of innate immune pathways, coupled with facultative growth outside tumours, constrains systemic dosing and narrows their therapeutic window. Clinical attenuation strategies, including modification of lipopolysaccharides, pathogenicity, metabolic auxotrophy, and invasiveness, have improved tolerability^7–9^, but expose a central tradeoff whereby attenuations compromise bacterial fitness and antitumour potency.

Synthetic biology offers a path to create dynamic gene circuits that sense environmental inputs and execute context-dependent responses^10–14^. Gene circuits such as quorum-controlled lysis systems, dynamic feedback controllers, and biocontainment kill switches allow precise regulation of growth, localization, and payload release^15–20^. As with oncolytic viruses and CAR-T cells, whose clinical realization required both the discovery of new engineering strategies and the integration of complementary features, bacterial therapies will require new innovations coupled with synthetic biology approaches that collectively enable control over safety, specificity, and therapeutic function.

Here, we establish an integrated engineering strategy within the probiotic *E. coli* Nissle 1917 (EcN), combining a newly identified lipopolysaccharide modification that reduces immunogenicity without compromising bacterial fitness, an AND-gate genetic circuit that restricts growth to the tumour microenvironment, and increased membrane permeability that enhances payload secretion while further improving safety **(Fig. 1a)**. This engineered strain expands the tolerable dose by up to 1,000-fold (route-dependent) across intraperitoneal, intravenous, and intracranial routes, drives durable complete tumour responses in colorectal carcinoma, and extends survival in both orthotopic glioblastoma and disseminated ovarian cancer. Together, these advances set the stage for broad translational applications.

**Fig. 1.**
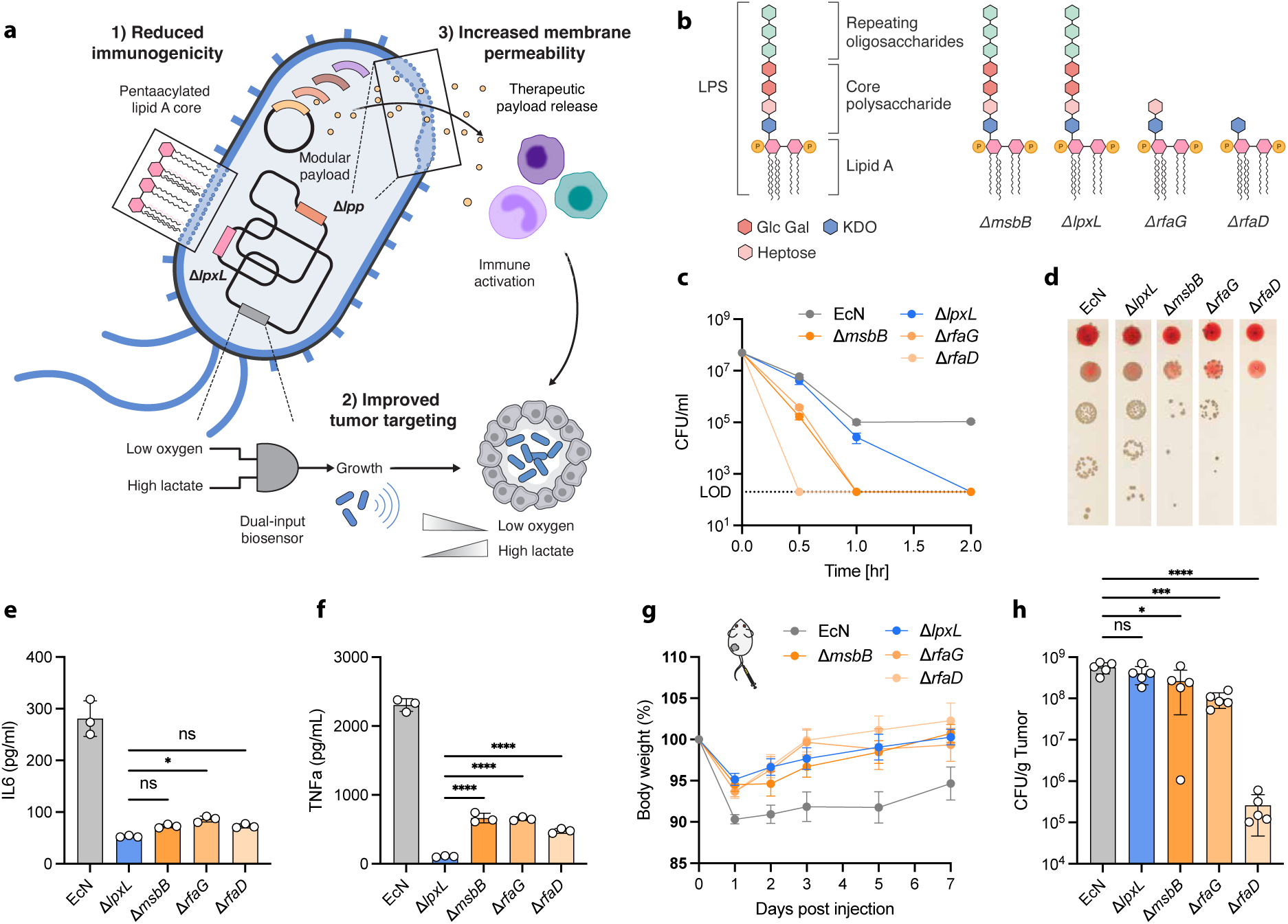
| Identification of lipopolysaccharide modifications that improve tolerability while preserving tumour colonization of EcN. **a**, Schematic of the engineered EcN platform incorporating Δ*lpxL* (reduced immunogenicity), Δ*lpp* (enhanced payload release), and a dual-input hypoxia/lactate biosensor (tumour-restricted growth). **b**, LPS structures of wild-type and mutant strains with deletions in *lpxL*, *msbB*, *rfaG*, or *rfaD*. **c**, Survival of strains in human whole blood over time. LOD, limit of detection. **d**, Representative LB agar plates after 30-minute blood incubation. **e**, **f**, IL-6 (**e**) and TNF-α (**f**) secretion by J774 macrophages stimulated with wild-type or LPS mutant strains (MOI = 0.1, 6 hours) (n = 3 biological replicates). **g**, Relative body weight of 4T1 tumour-bearing BALB/c mice after intravenous injection (10⁷ CFU) (n = 5 per group). **h**, Tumour colonization (CFU/g) on day 7 (n = 5 per group). Data are mean ± s.d. (**c**, **e**, **f**, **h**) or mean ± s.e.m. (**g**). Statistical significance by one-way ANOVA with Tukey’s multiple comparisons test (**e**, **f**) or Dunnett’s multiple comparisons test versus EcN (**h**); \**P* < 0.05, \*\*\**P* < 0.001, \*\*\*\**P* < 0.0001; ns, not significant.

## Decoupling immunogenicity from fitness

Lipopolysaccharide (LPS) is a dominant driver of innate immune activation, but its modification can limit tumour colonization^21^. We therefore screened a panel of LPS mutants in EcN to identify modifications that reduce immunogenicity without compromising tumour fitness. These mutations target two structurally distinct features of LPS: *msbB* and *lpxL* alter lipid A acylation, producing penta-acylated structures with reduced innate immune recognition, while *rfaG* and *rfaD* truncate the core oligosaccharide chain **(Fig. 1b)**. In vitro growth kinetics revealed that EcNΔ*lpxL* and EcNΔ*msbB* strains grew similarly to WT EcN, whereas EcNΔ*rfaG* and EcNΔ*rfaD* showed delayed growth, with combined deletions exhibiting compounded fitness costs **(Extended Data Fig. 1a, b)**. Because LPS also maintains outer membrane integrity and complement resistance, we next assessed blood survival across the panel. EcNΔ*lpxL* retained the highest viability in human whole blood over 2 hours, comparable to WT EcN at the 30-min timepoint, whereas EcNΔ*rfaD* and EcNΔ*rfaG* were rapidly cleared **(Fig. 1c, d)**. Combining deletions further exacerbated immune clearance, with both Δ*rfaG* Δ*msbB* and Δ*msbB* Δ*lpxL* being undetectable at 30 minutes **(Extended Data Fig. 1c)**. Given that LPS-mediated innate immune activation is a primary driver of systemic toxicity, we next measured inflammatory cytokine induction across the panel. All four mutants triggered substantially reduced IL-6 and TNF-α secretion from J774 murine macrophages compared to wild-type EcN **(Fig. 1e, f)**, with attenuation sustained across a broad range of MOIs and at 24 hours **(Extended Data Fig. 2a–d)**. This attenuation was confirmed in primary bone marrow-derived dendritic cells, where Δ*lpxL* consistently induced the lowest cytokine levels **(Extended Data Fig. 2e–h)**.

To evaluate the impact of LPS modifications on tolerability and tumour colonization in vivo, we administered these mutants intravenously in 4T1 triple-negative breast tumour-bearing mice. At an intravenous dose of 1×10⁷ colony-forming units (CFU), all mutants improved body weight retention relative to WT EcN, but differed markedly in their ability to maintain intratumoural colonization **(Fig. 1g)**. Notably, only EcN*ΔlpxL* maintained intratumoural bacterial burden at day 7 indistinguishable from WT EcN, whereas EcN*ΔrfaG*, EcN*ΔrfaD*, and EcN*ΔmsbB* strains showed significantly reduced tumour colonization **(Fig. 1h)**. At a higher dose of 2×10⁷ CFU, body weight retention was similarly improved across all mutant strains relative to WT EcN **(Extended Data Fig. 3a)**. Endotoxin activity, quantified by Limulus amebocyte lysate (LAL) assay, the clinical standard for pyrogen testing of injectable biologics, was substantially reduced in EcNΔ*lpxL* relative to WT EcN **(Extended Data Fig. 3b)**. Together, these data identify *lpxL* deletion as an attenuation strategy that minimizes pro-inflammatory signalling while preserving fitness and tumour colonization.

## Gating growth on hypoxia and lactate

Systemic bacterial therapy requires not only reduced immunogenicity but also active restriction of bacterial growth to tumours. Metabolic auxotrophy has been explored but has shown incomplete tumour colonization in clinical settings^7^. Synthetic circuits responsive to tumour hallmarks offer greater selectivity, and multiplexing these inputs through an AND gate further improves tumour specificity^22^. To impose tumour-restricted proliferation, we engineered a dual-input biosensor circuit coupling bacterial survival to hypoxia and elevated lactate **(Fig. 2a)**. We implemented this in a chromosomally integrated EcN*Δasd* auxotrophic chassis, in which bacterial growth depends on *asd* expression **(Extended Data Fig. 4a)**. *asd* encodes aspartate-semialdehyde dehydrogenase, which is essential for diaminopimelic acid (DAP) synthesis and peptidoglycan production. The *asd* gene was placed under a synthetic hypoxia– and lactate-responsive fusion promoter developed in this study, which combines binding sites for FNR, a transcription factor activated under low-oxygen conditions, and LldR, a lactate-responsive repressor whose binding is relieved by L-lactate. The two inputs form an AND gate that restricts growth to tumour-relevant conditions **(Fig. 2c)**. This design integrates both inputs through a single hybrid promoter controlling a single essential gene, creating a compact genetic architecture amenable to genomic integration while avoiding the variability, plasmid loss, and complexity of prior two-plasmid designs requiring multiple essential genes^22^.

**Fig. 2.**
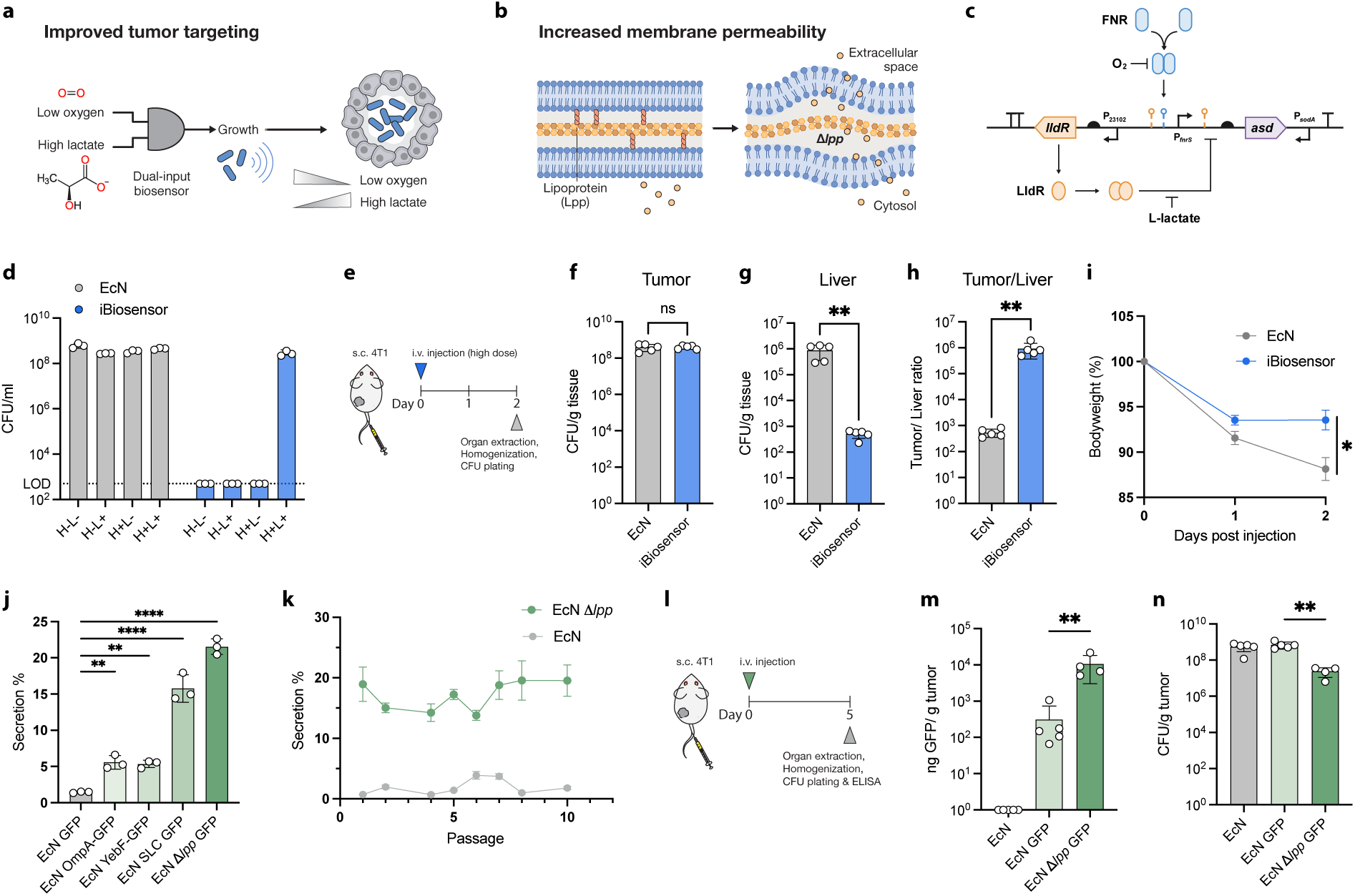
| A dual-input biosensor and *lpp* deletion enhance tumour specificity and enable efficient payload delivery. **a**, Schematic of the hypoxia/lactate AND-gate biosensor restricting bacterial growth to the tumour microenvironment. **b**, Schematic of Δ*lpp* deletion, which disrupts outer membrane anchoring and increases permeability for passive payload release. **c**, Genetic circuit diagram of the biosensor. FNR and LldR regulate expression of the essential gene *asd* under hypoxia and high lactate, respectively. **d**, In vitro CFU of wild-type EcN and the integrated biosensor (iBiosensor) strain after 24 hours of static culture across four combinations of hypoxia (H) and L-lactate (L) (n = 3 biological replicates). LOD, limit of detection. **e**, Experimental timeline for in vivo biodistribution following i.v. injection of wild-type EcN or the iBiosensor strain (2×10⁷ CFU) in 4T1 tumour-bearing mice. **f**–**h**, Bacterial colonization in the tumour (**f**) and liver (**g**), and the tumour-to-liver ratio (**h**), at day 2 (n = 5 per group). **i**, Body weight over the 2-day period (n = 5 per group). **j**, Percentage of GFP released by EcN expressing GFP alone (cytoplasmic), OmpA-tagged GFP, YebF-tagged GFP, GFP via the synchronized lysis circuit (SLC), or EcN Δ*lpp* expressing GFP (n = 3). **k**, GFP secretion efficiency over serial passages, confirming phenotypic stability (n = 4 for EcN Δ*lpp*, n = 3 for EcN). **l**, Experimental timeline for in vivo payload quantification. **m**, GFP accumulation (ng GFP/g tumour) in tumours at day 5 post-i.v. injection (n = 5, 5, and 4 for EcN, EcN GFP, and EcN Δ*lpp* GFP, respectively). **n**, Tumour colonization (CFU/g tumour) at day 5 (n = 5, 5, and 4 for EcN, EcN GFP, and EcN Δ*lpp* GFP, respectively). Data are mean ± s.d. (**d**, **f**–**h**, **j**, **m**, **n**) or mean ± s.e.m. (**i**, **k**). Statistical significance by Mann–Whitney test (**f**–**h**), two-way ANOVA with Šídák’s multiple comparisons test (**i**), or one-way ANOVA with Tukey’s multiple comparisons test (**j**, **m**, **n**); \**P* < 0.05, \*\**P* < 0.01, \*\*\**P* < 0.001, \*\*\*\**P* < 0.0001; ns, not significant.

Circuit performance was refined through rational design and iterative tuning of regulatory element strength and translation initiation. Tuning *lldR* expression and *asd* ribosome binding strength yielded a configuration supporting robust growth under hypoxic, lactate-rich conditions while suppressing proliferation in non-permissive environments **(Fig. 2d)**. In vivo, two days after a single intravenous injection **(Fig. 2e)**, the optimized biosensor strain (iBiosensor) reduced body weight loss **(Fig. 2i)**, maintained tumour colonization comparable to EcN in the 4T1 model **(Fig. 2f)** while reducing liver burden **(Fig. 2g)** and increasing the tumour-to-liver CFU ratio by >1,000-fold relative to WT EcN **(Fig. 2h)**. Off-target colonization was similarly reduced in independent biodistribution studies in the 4T1 model **(Extended Data Fig. 4b–d)** and in the MC38 model, where iBiosensor preserved tumour colonization while decreasing liver and spleen burden relative to WT EcN **(Extended Data Fig. 4e–g)**. Consistent with circuit-dependent growth, the EcN*Δasd* auxotrophic strain failed to colonize tumours **(Extended Data Fig. 4b–d)**. Together, these data demonstrate that the hypoxia-lactate biosensor restricts off-target proliferation without compromising tumour colonization across models.

## *lpp* deletion enhances payload release

Efficient bacterial therapy also requires extracellular release of therapeutic payloads. Existing strategies, including lysis circuits^15^ and engineered secretion systems^23,24^, can impose fitness costs, introduce genetic instability, or limit payload compatibility. As an alternative, we aimed to enable passive protein release by increasing outer membrane permeability through deletion of the *lpp* gene, which encodes Braun’s lipoprotein that anchors the outer membrane to peptidoglycan **(Fig. 2b)**^25–27^. Although *lpp* deletion has recently been used to render probiotics “leaky” for payload release, its efficiency relative to established secretion and lysis systems, and its stability over the prolonged timescales relevant to tumour colonization, remained undefined. EcN*Δlpp* maintained near-wild-type growth kinetics, indicating minimal fitness cost **(Extended Data Fig. 5a)**. Using GFP as a model payload of similar molecular weight to therapeutic cytokines, we compared Δ*lpp*-mediated release to several established secretion strategies: OmpA and YebF secretion tags^15,28,29^ and a synchronized lysis circuit (SLC). EcNΔ*lpp* released ∼20% of total GFP into the culture supernatant, compared with ∼5% for OmpA– and YebF-tagged GFP and ∼15% for the SLC **(Fig. 2j)**. This secretion phenotype was genetically stable, persisting at consistent levels over 10 serial passages of continuous subculturing, a property essential for bacteria that must maintain function during extended tumour colonization **(Fig. 2k)**. To confirm that enhanced secretion translated to improved payload delivery *in vivo*, we administered GFP-expressing strains intravenously to 4T1 tumour-bearing mice and quantified intratumoural GFP at day 5 **(Fig. 2l)**. The EcNΔ*lpp* strain yielded >30-fold higher GFP accumulation per gram of tumour than the WT GFP-expressing strain, despite a slight reduction in bacterial colonization levels, confirming that the enhanced membrane permeability phenotype drives superior payload delivery in the tumour **(Fig. 2m, n)**.

## An expanded therapeutic window

We consolidated all three modifications into a single strain, BLL (Biosensor, Δ*lpxL*, Δ*lpp*), and assessed whether the combined modifications produced improvements in tolerability, with each modification contributing independently. Each component contributes to tumour selectivity through a distinct mechanism: the biosensor restricts growth to environments with low oxygen and high lactate, while Δ*lpxL* and Δ*lpp* act systemically: Δ*lpxL* attenuates host innate immune activation, while Δ*lpp* increases outer membrane permeability, rendering bacteria more susceptible to clearance in non-tumour tissues outside the tumour microenvironment. In a biodistribution study comparing individual modifications and the combined BLL strain in 4T1 tumour-bearing BALB/c mice, all strains colonized tumours at comparable levels, though EcNΔ*lpp* and BLL showed modestly reduced intratumoural titers consistent with the increased membrane permeability resulting from Δ*lpp* **(Extended Data Fig. 5b)**. Off-target colonization reflected the independent contribution of each modification: iBiosensor, Δ*lpxL*, and Δ*lpp* each partially reduced liver and spleen burden relative to wild-type EcN, while BLL, combining all three, achieved the greatest reduction in both organs **(Extended Data Fig. 5c, d)**. Body weight loss followed the same trend, with each individual modification conferring partial protection and BLL showing the least toxicity, consistent with the convergence of metabolic containment and immune attenuation acting through independent mechanisms **(Extended Data Fig. 5e)**.

Serum cytokine profiling 4 hours after a single intravenous (i.v.) injection revealed that BLL broadly suppressed the induction of pro-inflammatory mediators relative to wild-type EcN, with levels approaching those of PBS-treated controls **(Fig. 3a)**. BLL-treated mice showed lower serum levels of TNF-α, IL-6, IL-1β, GROα/CXCL1, MIP-2α/CXCL2, and G-CSF **(Fig. 3b–g)**. This broad suppression across multiple cytokine axes, spanning both alarmin (IL-1β) and chemokine (GROα, MIP-2α) pathways, indicates that the combined modifications dampen host innate immune recognition at multiple levels rather than attenuating a single signalling axis.

**Fig. 3.**
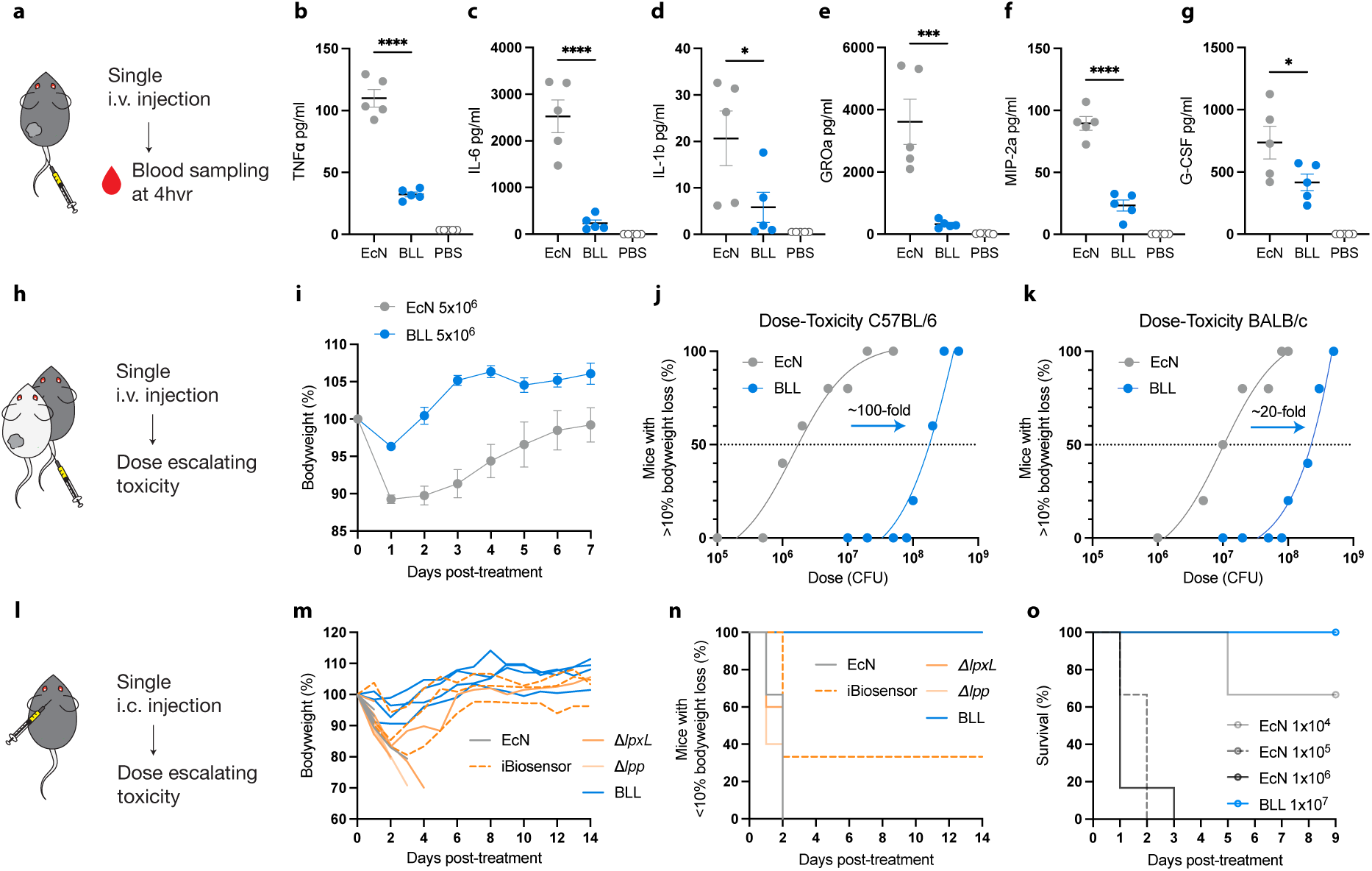
| BLL engineering dramatically improves tolerability across administration routes and mouse strains. **a**, Schematic of serum cytokine profiling 4 hours after a single i.v. injection. **b**–**g**, Serum levels of TNF-α (**b**), IL-6 (**c**), IL-1β (**d**), GROα/CXCL1 (**e**), MIP-2α/CXCL2 (**f**), and G-CSF (**g**) in EcN-, BLL-, or PBS-treated mice, showing broadly reduced pro-inflammatory cytokine and chemokine induction in BLL-treated mice (n = 5 per group). **h**, Schematic of i.v. dose-escalation toxicity study. **i**, Relative body weight following i.v. injection of EcN or BLL at 5×10⁶ CFU (n = 5 per group). **j**, **k**, Dose-toxicity curves showing the fraction of mice with >10% body weight loss in C57BL/6 (**j**) and BALB/c (**k**) mice, revealing ∼100-fold and ∼20-fold increases in maximum tolerated dose for BLL, respectively (n = 5 to 10 mice per dose). **l**, Schematic of intracranial (i.c.) dose-escalation study in healthy mice. **m**, Individual body weight traces and **n**, freedom from >10% body weight loss in healthy C57BL/6 mice following i.c. injection of EcN, iBiosensor, Δ*lpxL*, Δ*lpp*, or BLL (1×10⁶ CFU; n = 6, 3, 5, 5, and 4, respectively). In n, mice were scored as events on the first day of >10% weight loss from baseline or death, and survivors were censored at day 14. **o**, Kaplan–Meier survival curves after i.c. injection, demonstrating that BLL is well tolerated at 1×10⁷ CFU whereas EcN caused lethality at all doses tested (≥1×10⁴ CFU), with complete lethality at ≥1×10⁵ CFU (n = 3, 3, and 6 for EcN at 1×10⁴, 1×10⁵, and 1×10⁶ CFU, respectively, and n = 5 for BLL at 1×10⁷ CFU). Data are mean ± s.e.m. (**b**–**g**, **i**). Statistical significance determined by one-way ANOVA with Dunnett’s multiple comparisons test versus EcN (**b**–**g**) or log-rank test (**n**, **o**); \**P* < 0.05, \*\*\**P* < 0.001, \*\*\*\**P* < 0.0001.

We next performed dose-escalation toxicity studies to quantify the therapeutic window expansion. BLL-treated mice showed improved body weight trajectories compared to EcN-treated mice in MC38 tumour-bearing C57BL/6 mice **(Fig. 3h, i)**. Dose-toxicity curves revealed that the maximum tolerated dose (MTD) of BLL was approximately 100-fold higher than that of EcN in C57BL/6 mice **(Fig. 3j)**. This increased therapeutic window was also observed in BALB/c mice with BLL exhibiting an approximately 20-fold higher MTD than EcN **(Fig. 3k)**. Across the dose range tested, BLL-treated mice consistently retained more body weight than EcN-treated mice, and BLL maintained comparable intratumoural colonization regardless of dose in both mouse backgrounds **(Extended Data Fig. 6a–h)**.

We also evaluated BLL tolerability following intracranial (i.c.) administration, a route relevant to central nervous system (CNS) malignancies where direct injection bypasses the blood-brain barrier but the brain’s limited tolerance to neuroinflammation imposes strict safety constraints **(Fig. 3l)**. A study comparing individual modifications at matched i.c. doses showed that tolerability required the full combination: EcN, iBiosensor, ΔlpxL, and Δlpp each caused >10% body weight loss or lethality, whereas no BLL-treated animal exceeded 10% body weight loss **(Fig. 3m, n)**. This indicates that integrating all three modifications provides a safety margin beyond any individual engineering component. Consistent with this margin, i.c. injection of wild-type EcN caused dose-dependent lethality, with mortality occurring even at the lowest dose tested (1×10⁴ CFU), whereas BLL was tolerated at 1×10⁷ CFU, a 1000-fold higher dose, with all mice surviving the entire 9-day observation period **(Fig. 3o, Extended Data Fig. 7a, b)**.

To assess whether this expanded therapeutic window extended to intraperitoneal (i.p.) delivery, we tested BLL in a disseminated ovarian cancer model. Mice were challenged i.p. with mFTC-ΔTD-luc cells, followed by i.p. bacterial treatment. Whereas EcN caused treatment-related mortality at its tested dose, BLL was tolerated with no deaths at a 10-fold higher dose of bacteria **(Extended Data Fig. 8a, b)**. Together, these data establish a substantial improvement in the tolerated dose across i.v., i.c., and i.p. administration routes and demonstrate that the integrated engineering strategy produces a safety profile that exceeds that of any individual modification alone.

## Efficacy across tumour types and routes

Having established that BLL expands the therapeutic window, we next evaluated whether this improvement translates to antitumour efficacy across distinct tumour contexts and delivery routes.

We first tested BLL in an orthotopic syngeneic GL261 glioblastoma model. Mice received i.c. GL261 engraftment followed by i.c. BLL injection two weeks later **(Fig. 4a)**. To model co-administration with standard-of-care therapy, a subset of mice received two cycles of daily intraperitoneal temozolomide (TMZ). TMZ alone extended median survival from 4 days (PBS) to 8 days, while BLL monotherapy extended median survival to 9 days. The combination of i.c. BLL with systemic TMZ further extended median survival to 14 days, exceeding either monotherapy alone **(Fig. 4b)**. This enhanced effect is consistent with mechanistic complementarity: BLL localizes to tumour and peritumoural tissue and induces immunostimulatory responses, whereas TMZ targets proliferating tumour cells as an alkylating prodrug^30,31^. Immunohistochemistry of brain sections from BLL-treated mice confirmed that bacterial biodistribution, detected by LPS staining, was restricted to the peritumoural region, with no detectable bacterial burden in surrounding healthy brain parenchyma **(Fig. 4c)**.

**Fig. 4.**
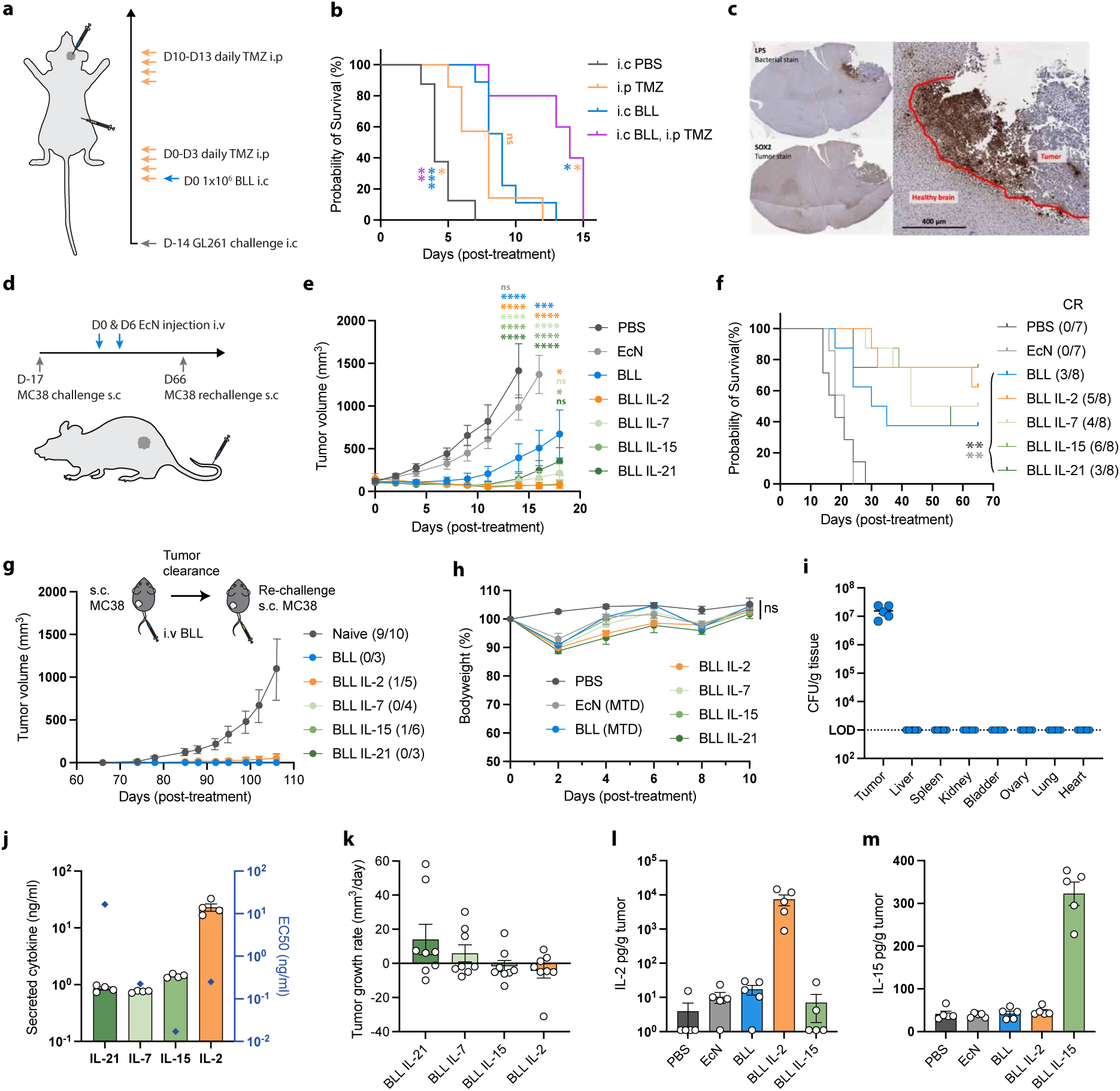
| BLL demonstrates therapeutic efficacy in orthotopic glioblastoma and subcutaneous colorectal tumour models. **a**, Schematic of the orthotopic GL261 study. Mice received i.c. GL261 engraftment on day −14, i.c. BLL (1×10⁶ CFU) on day 0, and i.p. temozolomide (TMZ, 1 mg) daily on days 0 to 3 and 10 to 13. **b**, Kaplan–Meier survival (n = 8, 7, 9, and 5 for i.c. PBS, i.p. TMZ, i.c. BLL, and i.c. BLL + i.p. TMZ). Asterisks are coloured to match the corresponding group and denote its pairwise log-rank comparison with the adjacent survival curve. **c**, Representative LPS (bacteria, top) and SOX2 (tumour, bottom) immunohistochemistry of a BLL-treated GL261 brain section at endpoint. Red outline demarcates the boundary between the tumour and the healthy brain. Scale bar, 400 µm. **d**, Schematic of the subcutaneous MC38 study. Mice received the MC38 challenge on day −17, i.v. treatment on days 0 and 6 at the maximum tolerated dose for each strain (EcN, 2×10^6^ CFU; BLL and cytokine-expressing BLL strains, 2×10^8^ CFU per injection), and MC38 rechallenge on day 66. **e**, Mean tumour volume over time (n = 7, 7, 8, 8, 8, 8, and 8 for PBS, EcN, BLL, BLL-IL2, BLL-IL7, BLL-IL15, and BLL-IL21). Asterisk colour indicates the tested group, following the same colour scheme as the group labels. At days 14, 16, and 18, groups were compared against PBS, EcN, and BLL, respectively; the comparator advanced as earlier control groups reached humane endpoints. **f**, Kaplan–Meier survival; complete response (CR) fractions are shown next to each group label. Asterisk colour indicates the comparator control. **g**, Tumour volume following MC38 rechallenge (1×10⁶ cells) of complete responders versus naïve age-matched controls. Fractions indicate the number of mice that developed a tumour out of the total rechallenged. **h**, Relative body weight following i.v. treatment with PBS, EcN, BLL, or each cytokine-armed strain at the maximum tolerated dose (n as in E). **i**, Biodistribution of BLL (CFU/g tissue) at day 10 post-treatment across the tumour and organs (n = 5). Dotted line, limit of detection (LOD). **j**, In vitro cytokine payload activity. Left axis (bars): concentration of each cytokine (IL-21, IL-7, IL-15, IL-2) secreted by the corresponding cytokine-armed BLL strain into bacterial culture supernatant (ng/ml; n = 4). Right axis (diamonds): the concentration required for half-maximal activity (EC50; see Methods). **k**, Tumour growth rate (mm³/day) for each cytokine-armed strain, calculated for each animal as the mean of the volume changes between consecutive timepoints (ΔV/Δt) over the treatment course (circles, individual mice; n = 8 per group). **l**, **m**, Intratumoural IL-2 (**l**) and IL-15 (**m**) concentration (pg/g tumour) at day 2 post-treatment (circles, individual mice; n = 5 per group). Data are mean ± s.e.m. (**e**, **g**–**m**). Statistical significance by log-rank test (**b**, **f**) or one-way ANOVA with Dunnett’s multiple comparisons test at days 14, 16, and 18 (**e**); \**P* < 0.05, \*\**P* < 0.01, \*\*\**P* < 0.001, \*\*\*\**P* < 0.0001.

We next evaluated BLL as a systemic delivery vector in a subcutaneous MC38 colorectal tumour model (**Fig. 4d**). The modular architecture of BLL enables expression of diverse therapeutic payloads, and we engineered strains expressing one of four γ-chain cytokines (IL-2, IL-7, IL-15, IL-21). The IL-15 payload was delivered as an IL-15Rα sushi domain–IL-15 fusion (an RLI-type superagonist)^32^, hereafter referred to as IL-15. MC38-bearing mice received two intravenous injections of PBS, wild-type EcN, BLL, or cytokine-expressing BLL strains, on treatment days 0 and 6, starting on day 17 post-engraftment. Bacterial strains were administered at their respective MTDs. Relative to PBS and EcN controls, all BLL-treated strains suppressed tumour growth **(Fig. 4e)** and extended survival, with BLL-IL2 and BLL-IL15 conferring the greatest benefit **(Fig. 4f, Extended Data Fig. 9)**. BLL alone achieved complete responses in 3 of 8 mice, while cytokine arming further improved this to 5 of 8 (BLL-IL2) and 6 of 8 (BLL-IL15), compared to 0 of 7 in PBS and EcN controls.

Mice that completely rejected tumours were rechallenged with MC38 cells without further treatment, and nearly all remained tumour-free, consistent with durable antitumour immunity **(Fig. 4g).** BLL strains were well tolerated, exhibiting body weight changes comparable to wild-type EcN despite administration at a substantially higher dose **(Fig. 4h).** Biodistribution analysis confirmed tumour-restricted colonization, with bacteria undetectable in off-target organs **(Fig. 4i)**. We next compared therapeutic efficacy with in vitro cytokine activity. While cytokine secretion alone did not account for differences in efficacy, tumour growth inhibition was more closely associated with cytokine expression relative to its effective concentration (EC50), with payloads exceeding their EC50 generally producing greater tumour suppression **(Fig. 4j, k)**. Intratumoural ELISA of the two highest-expressing payloads, IL-2 and IL-15, confirmed in vivo cytokine expression consistent with their relative in vitro secretion levels **(Fig. 4l, m)**. Together, these findings suggest that combining secretion level with intrinsic cytokine potency may provide a useful framework for prioritizing payloads and predicting in vivo efficacy.

To assess generalizability to additional tumour types and genetic backgrounds, we tested BLL-IL2 in BALB/c mice bearing orthotopic 4T1 mammary tumours. BLL-IL2 reduced tumour growth relative to PBS and BLL controls, with elevated intratumoural IL-2 confirmed by ELISA **(Extended Data Fig. 10a–d)**. We also evaluated BLL in the disseminated ovarian cancer model, where i.p.-delivered BLL reduced tumour progression measured by bioluminescence imaging and maintained intratumoural colonization at endpoint **(Extended Data Fig. 8c–e)**. With intravenous delivery, BLL-IL15 significantly extended median survival compared to PBS **(Extended Data Fig. 8f)**. Together, these data demonstrate that the BLL platform converts the expanded therapeutic window into a therapeutic advantage across multiple tumour types, anatomical contexts, and administration routes.

## Discussion

The narrow therapeutic window imposed by dose-limiting toxicity remains the central barrier to clinical translation of bacterial cancer therapies. Prior efforts have addressed individual aspects of this problem, including attenuation, tumour targeting, and payload delivery, but these strategies implemented in isolation have not sufficiently expanded the therapeutic window. Here, we show that integrating three mechanistically independent modifications into a single EcN strain substantially improves safety, expanding the maximum tolerated dose across multiple delivery routes and enabling robust antitumour efficacy across distinct tumour models. Beyond increasing tolerable dosing, the expanded therapeutic window enables interrogation of BLL across a broad dose range, providing an opportunity to analyse dose-response relationships and mechanisms inaccessible with wild-type EcN. These gains were observed in both C57BL/6 and BALB/c mice, which exhibit divergent immune polarization, indicating that the benefit is not restricted to a single host background. Importantly, because mice are relatively resistant to endotoxin compared with humans^33,34^, the magnitude of the safety improvement may be underestimated in our models and warrants direct evaluation in higher species.

Each component of BLL addresses a distinct biological constraint and illustrates a broader design principle for microbial therapeutics. The LPS mutant screen that guided selection of *lpxL* deletion demonstrates that reduced immunogenicity alone is insufficient as a criterion for attenuation; modifications must also preserve the bacterial fitness required for therapeutic function. Although all four LPS modifications reduced macrophage cytokine induction comparably, only EcN Δ*lpxL* maintained blood survival and intratumoural colonization near wild-type levels, demonstrating that immunogenicity and fitness can be genetically decoupled rather than necessarily traded against one another. However, even bacteria with reduced immunogenicity can proliferate in off-target tissues, amplifying otherwise attenuated inflammatory signals and producing dose-limiting toxicity. A complementary strategy is therefore needed to restrict where bacterial growth can occur. Metabolic auxotrophies have historically provided one approach to containment, but their dependence on specific metabolites can constrain bacterial fitness and make tumour growth contingent on nutrient availability. In contrast, genetically encoded biosensors can couple bacterial fitness to broadly conserved features of tumour physiology rather than the availability of a single metabolite. Our dual-input “AND” gate integrates hypoxia and lactate, effectively converting a facultative bacterium into a synthetic “tumorobe” whose growth is preferentially coupled to the tumour microenvironment. Chromosomal integration further provides genetic stability without dependence on plasmid maintenance, reducing opportunities for plasmid loss and horizontal gene transfer during prolonged colonization. Finally, EcN Δ*lpp* addresses the distinct challenge of payload delivery from Gram-negative bacteria by increasing membrane permeability for passive protein release while simultaneously enhancing clearance from off-target tissues. EcN Δ*lpp* also reduces TLR2 stimulation and increases susceptibility to antibiotics such as vancomycin, providing additional features favourable for clinical translation^35,36^. Together, these modules illustrate how orthogonal engineering strategies can be layered to independently control immunogenicity, spatial growth, and therapeutic delivery rather than relying on attenuation alone.

The efficacy data demonstrate that the expanded therapeutic window translates directly into antitumour activity. BLL alone achieved durable complete responses in the MC38 model, while arming BLL with γ-chain cytokines further improved response rates, with IL-2 and IL-15 showing the greatest benefit. The modular platform enables rapid evaluation of alternative or combination payloads across tumour contexts, and our data suggest that in vitro levels may provide an initial indicator of therapeutic activity in vivo. Notably, IL-2 and IL-15 were produced at concentrations substantially exceeding their reported EC50 values, which may contribute to their greater efficacy relative to the other cytokines tested. Efficacy in orthotopic glioblastoma and disseminated ovarian cancer models, including compatibility with temozolomide, further supports the breadth of the platform across tumour types and treatment contexts. Several limitations of the current study should be noted. The mechanisms underlying antitumour responses to individual payloads have not been fully characterized, and the current plasmid-based payload systems may require chromosomal integration or essential gene complementation for future clinical development. Together, these findings establish BLL as a modular therapeutic platform and demonstrate how the integration of safety, containment, and delivery can transform bacteria into broadly applicable living therapeutics.

## Methods

### Mice

All animal experiments were approved by the Institutional Animal Care and Use Committee (Columbia University, protocol AACF5755). Female BALB/c and C57BL/6 mice (Jackson Laboratories), 7–8 weeks old, were maintained in accordance with all rules for animal research at Columbia University. Mice were housed in a facility with a 12 h light–dark cycle and provided unrestricted access to food and water. The housing facility was maintained at 21–24 °C and 40– 60% humidity. Sample sizes were determined on the basis of previous studies and pilot experiments. Dose-toxicity curves were defined by the fraction of mice experiencing greater than 10% body weight loss, with the maximum tolerated dose (MTD) defined as the dose at which 50% of mice experience 10% body weight loss.

### Cell lines

CT26 and 4T1 cells were maintained in RPMI supplemented with 10% (vol/vol) fetal bovine serum (FBS), 1% (vol/vol) MEM non-essential amino acids solution, and 100 U ml⁻¹ penicillin-streptomycin (Pen/Strep). J774, MC38, and GL261 cells were maintained in Dulbecco’s modified Eagle’s medium (DMEM) supplemented with 10% (vol/vol) FBS, 1× GlutaMax, 1% (vol/vol) MEM non-essential amino acids solution, and 100 U ml⁻¹ penicillin-streptomycin. mFTC cells were maintained in MEM alpha containing L-glutamine, ribonucleosides, and deoxynucleosides, supplemented with 10% (vol/vol) FBS, 10 µg ml⁻¹ insulin-transferrin-sodium selenite, and 0.02 µg ml⁻¹ beta-estradiol. Cultures were maintained in a humidified 37 °C, 5% CO₂ incubator. Cells were confirmed mycoplasma-free.

### Bacterial strains and plasmid construction

Genomic deletions were generated using a CRISPR-Cas9–assisted λ-Red recombineering system^37^. LPS synthesis genes (*msbB*, *lpxL*, *rfaG*, and *rfaD*), the essential gene *asd*, and the lipoprotein gene *lpp* were targeted for deletion. For each knockout, a crRNA targeting the gene of interest, together with 500 bp homology arms flanking the target, was cloned into the pTargetF backbone. Bacteria were first transformed with the pCas plasmid, which encodes Cas9 for counterselection and the λ-Red recombinase for homologous recombination; the corresponding pTargetF plasmid was then introduced to drive the deletion. For biosensor integration, circuit components were assembled into a CRIM (pAH162) backbone carrying a tetracycline resistance gene by Gibson assembly and integrated site-specifically at the φ80 locus^38^.

Cytokine and reporter payloads (GFP, IL-2, IL-7, IL-15, and IL-21) were expressed from an axe/txe toxin-antitoxin–stabilized plasmid backbone carrying a p15A origin of replication and a kanamycin resistance gene^39^. Each payload coding sequence was ordered as a gene fragment (Twist Bioscience) and inserted into the backbone by Gibson assembly. Expression was driven by the constitutive J23100 promoter (Anderson promoter collection), and ribosome binding sites were designed using the RBS Calculator^40^. All knockouts, integrations, and plasmid constructs were verified by PCR and whole-plasmid sequencing. Nucleotide sequences of the cytokine payload expression cassettes are provided in Supplementary Table 1.

### Blood bactericidal assay

LPS and *lpp* mutant bacterial strains were cultured overnight in LB broth. Cultures were pelleted by centrifugation at 3,000 × g for 10 minutes, resuspended in PBS, and adjusted to an OD₆₀₀ of 5. For each strain, 150 µl of human whole blood from a single donor (Innovative Research) was aliquoted into three wells of a 96-well plate, and 1.5 µl of bacterial suspension was added to each well and incubated at 37 °C. At designated time points, serial dilutions of each sample were prepared in PBS and plated on LB agar supplemented with erythromycin, then incubated overnight at 37 °C. Colony-forming units (CFU) were counted the following day.

### Dendritic cell and macrophage stimulation assay

Bone marrow-derived dendritic cells (BMDCs) were generated from the bone marrow of C57BL/6 mice and differentiated in RPMI-1640 supplemented with 10% (vol/vol) FBS, 1% penicillin-streptomycin, and 20 ng ml⁻¹ GM-CSF for 7 days, with medium replenished every 2 to 3 days^41^. BMDCs or J774 cells were seeded in 96-well plates at 5×10⁵ cells per well and allowed to adhere overnight. LPS mutant *E. coli* strains were grown for 18 hours in LB broth at 37 °C with shaking. Live bacteria were collected by centrifugation at 3,000 × g for 10 minutes, washed twice with PBS, resuspended in PBS, and added to BMDCs or J774 cells at a multiplicity of infection (MOI) ranging from 0.01 to 100. Control wells received PBS alone. Cells were incubated for 6 or 24 hours, after which supernatants were collected for cytokine quantification by ELISA. Mouse IL-6 and TNF-α ELISAs were performed using DuoSet ELISA kits (R&D Systems) according to the manufacturer’s protocols. All experiments were performed in biological triplicate.

### Endotoxin quantification

Endotoxin activity of wild-type EcN, the LPS-attenuated EcN Δ*lpxL* and Δ*msbB* strains, and a ClearColi low-endotoxin reference was quantified using a *Limulus* amebocyte lysate (LAL) chromogenic endpoint assay (Hycult Biotech, cat. no. HIT302). Whole bacteria were assayed at 10³ CFU per well, and endotoxin activity was determined against a standard curve of known endotoxin concentrations according to the manufacturer’s instructions.

### Characterization of biosensor-coupled growth in vitro

Each bacterial strain was grown overnight in liquid LB, then subcultured overnight in freshly reconstituted 1× MOPS EZ Rich defined medium supplemented with 100 µg ml⁻¹ DAP. Overnight cultures grown in MOPS EZ Rich medium were washed three times in PBS before inoculation into 4 ml of MOPS EZ Rich medium with or without L-lactate (40 mM). Hypoxia was established for these static liquid cultures using a BD GasPak EZ anaerobic chamber with sachets. All conditions were inoculated with 10⁵ CFU of bacteria and grown statically for 24 hours in a 37 °C incubator, after which each culture was serially diluted in triplicate and plated on LB agar containing antibiotics and DAP for CFU enumeration.

### Bacterial biodistribution in vivo

To assess bacterial colonization in vivo, bacterial strains were cultured overnight in LB broth at 37 °C with shaking. Cultures were pelleted by centrifugation at 3,000 × g for 10 minutes, washed three times, and resuspended in PBS. Mice were administered bacteria via tail-vein injection at a defined dose. At designated time points post-administration, mice were euthanized, and tissues of interest (tumour, liver, and spleen) were harvested under sterile conditions. Tissues were weighed, homogenized in PBS using a tissue homogenizer, and serially diluted. Diluted homogenates were plated on LB agar supplemented with the appropriate antibiotics for strain selection and incubated overnight at 37 °C. Colony-forming units (CFU) were counted the following day and normalized to tissue weight to quantify bacterial burden.

### Protein expression and secretion assay

To evaluate protein expression and secretion, wild-type and Δ*lpp* mutant bacterial strains transformed with GFP or cytokine payloads were grown for 18 hours in LB broth at 37 °C with shaking. Cultures were centrifuged at 3,000 × g for 10 minutes to separate the supernatant from the bacterial pellet. For GFP-expressing strains, fluorescence was measured directly in the supernatant and in the resuspended pellet using a plate reader. For cytokine-expressing strains, the supernatant was collected, and the pellet was resuspended in PBS and lysed by sonication, followed by centrifugation to remove cell debris; cytokine levels in the supernatant and lysate were quantified using DuoSet ELISA kits (R&D Systems) according to the manufacturer’s protocols. Secretion efficiency was calculated as supernatant signal divided by total signal (supernatant plus pellet), expressed as a percentage. For the passage stability study, overnight cultures were diluted 1,000-fold daily to establish subcultures over 10 days, with protein expression and secretion measured by the same method at each passage.

### Serum cytokine and chemokine profiling

Mice received a single intravenous injection of PBS, wild-type EcN, or an engineered strain, and whole blood was collected 4 hours after injection. Serum was separated by centrifugation and stored at −80 °C until analysis. Serum concentrations of cytokines and chemokines were measured using the ProcartaPlex Mouse Cytokine & Chemokine Panel 1A 36-plex (Invitrogen/Thermo Fisher Scientific, catalogue no. EPX360-26092-901), a Luminex xMAP bead-based multiplex immunoassay, according to the manufacturer’s instructions. In brief, 25 µl of serum or cytokine standard was incubated with antibody-conjugated magnetic capture beads, followed by biotinylated detection antibodies and streptavidin– phycoerythrin, with washes between steps. Beads were acquired on a Luminex 200 instrument, and analyte concentrations were interpolated from analyte-specific standard curves.

### Protein quantification in vivo

Tumours were harvested from treated mice, weighed, and homogenized in PBS using a tissue homogenizer. Tumour homogenates were centrifuged at 3,000 × g for 10 minutes, and GFP or cytokine levels in the supernatant were quantified using an Abcam GFP ELISA kit and DuoSet ELISA kits (R&D Systems), respectively. Protein levels were normalized to tumour weight to assess bacterial protein secretion within the tumour microenvironment.

### Colorectal and mammary tumour inoculation and treatment in vivo

To establish tumours in vivo, MC38 (colorectal) or 4T1 (mammary) tumour cells were cultured as described above. Cells were harvested at 80–90% confluency, washed with PBS, and resuspended in PBS. MC38 cells were implanted into C57BL/6 mice, and 4T1 cells into BALB/c mice, by subcutaneous injection of 1×10⁶ cells in 100 µl PBS into the right flank using a 27G needle. For the orthotopic mammary model, 4T1 cells were instead inoculated directly into the mammary fat pad (MFP) of BALB/c mice by injection of 5×10⁵ cells in 50 µl PBS using a 27G needle. Tumour growth was monitored by caliper measurements, and tumour volume was calculated using the formula V = ½ × length × width², with length being the larger diameter.

Once tumours reached ∼100 mm³, mice were randomly assigned to treatment groups. Bacterial strains were cultured overnight in LB broth, centrifuged at 3,000 × g for 10 minutes, washed three times, and resuspended in PBS. Mice were injected with bacteria via the tail vein at a defined dose in 100 µl PBS. Control groups received PBS alone. Mice were monitored daily for tumour growth, body weight, and clinical signs of toxicity. At designated time points, tumours were harvested for quantification of bacterial burden (CFU counts) and for cytokine or protein expression analysis. Tumour tissues were weighed, homogenized in PBS, and processed for CFU plating or ELISA measurement. All experiments were performed in compliance with institutional animal care guidelines.

To assess protective immunity, mice that achieved complete tumour regression following treatment were rechallenged with a second subcutaneous injection of 1×10⁶ MC38 cells in 100 µl PBS into the contralateral flank using a 27G needle. Naïve mice were inoculated in parallel as controls. Tumour growth was monitored by caliper measurements as described above, and the fraction of mice remaining tumour-free was recorded.

### Glioblastoma tumour inoculation and treatment in vivo

Female 12–14-week-old C57BL/6 mice were anaesthetized with 1–2% isoflurane (Covetrus) and 2.5% oxygen. Mice were shaved, and depilatory cream (Veet) was applied for 2 minutes and then wiped off. Mice received subcutaneous injections of 100 µl electrolyte solution with 10% dextrose (Covetrus) in the left flank, 3.25 mg kg⁻¹ buprenorphine extended-release (Covetrus) in the right flank, and 100 µl of 2% lidocaine solution (Covetrus) in the scalp. The head was secured in the nose cone of a World Precision Instruments stereotactic frame, and the ear bars were fixed so that the head was flat and secure. The site was cleaned three times with iodine and isopropyl alcohol prep pads (Fisher), alternating between pad types. The scalp was opened with a 1 cm incision, and a 10 µl gastight Hamilton syringe fitted with a 26-gauge needle attached to the stereotactic frame was aligned with the bregma. A burr hole was drilled 2 mm to the right and 2 mm posterior to the bregma using a 1 mm bit on an Ideal Micro-Drill (Cell Point Scientific). Either 7 µl of a GL261 suspension (1.6×10⁷ cells ml⁻¹) or a bacterial suspension was loaded into the needle. The needle was advanced 4 mm into the brain and retracted 0.5 mm to a final depth of 3.5 mm, and 5 µl of the sample was injected over 5 minutes. The needle was withdrawn 1 mm every 2 minutes. The burr hole was sealed with bone wax, and the incision was sutured (Oasis PGA sutures, Fisher Scientific) and sealed with 3M Vetbond tissue adhesive. Mice were monitored daily for two days for post-operative checks of the incision and body weight. Mice were treated two weeks after grafting.

### Immunohistochemistry

Immunohistochemistry was performed on 4% paraformaldehyde-fixed, paraffin-embedded tissue sections (5 µm thick) to analyse LPS and SOX2 in bacteria-treated mouse brains. Sections were deparaffinized in xylene and rehydrated through 100% ethanol, 95% ethanol (2 × 5 minutes), and 75% ethanol (1 × 5 minutes). Slides were washed in water, and antigen retrieval was performed in 10 mM citrate buffer (pH 6) in a pressure cooker for 10 minutes. After a 30-minute cooling period, slides were washed in PBS (pH 7.4), and sections were incubated in primary antibody solutions overnight at room temperature. Staining was visualized using the VECTASTAIN ABC kit (Vector Laboratories) and DAB chromogen. The following primary antibodies were used: rabbit anti-SOX2 (Cell Signaling Technology, 1:1000) and mouse anti-LPS (Hycult Biotech, 1:1000). Immunoperoxidase-stained sections were scanned at 40× using a Leica SCN400 digital slide scanner. SOX2 and LPS staining were classified as positive or negative based on immunoperoxidase reactivity and quantified using QuPath.

### Temozolomide chemotherapy administration

Temozolomide (Millipore Sigma, PHR1437) was dissolved in DMSO to make a 15 mg ml⁻¹ stock solution, which was diluted with PBS to 1 mg ml⁻¹ and sterile-filtered through a 0.22 µm filter. When treated on the same day as intracranial bacterial injection, grafted C57BL/6 mice were allowed to recover until ambulatory before dosing. Mice were injected intraperitoneally with 1 ml of the 1 mg ml⁻¹ temozolomide solution (1 mg per mouse) using a 26-gauge beveled needle.

### Disseminated ovarian tumour model and treatment in vivo

Female albino C57BL/6 mice were intraperitoneally injected with 10⁶ mFTC-ΔTD-luc cells on day 0 to establish disseminated tumour nodules within the peritoneal cavity. Tumour engraftment was monitored by cell luminescence using an In Vivo Imaging System (IVIS). Saline or bacterial groups were then administered either intraperitoneally or intravenously in separate cohorts. Mice were monitored periodically for body condition score, body weight, bacterial colonization (by IVIS), and tumour growth until endpoint. At the endpoint, tumours, uterus, ovaries, and pancreas were imaged for bacterial and cancer cell luminescence and processed for bacterial colony quantification as described above.

The parental mFTC-Δ cells were generated in the Drapkin laboratory from mouse oviduct secretory epithelial cells (mFTC) by CRISPR-mediated knockout of *Trp53*, *Pten*, and *Brca2*. The mFTC-ΔTD line was then established from tumour-derived cells isolated after implantation of mFTC-Δ cells into mice. This line was subsequently transduced with lentiviral luciferase (pGreenFire1-ISRE) to constitutively express the reporter for live imaging.

### Estimation of functional payload activity

To place secretion levels on a common functional scale, we compared each cytokine’s in vitro secreted amount with its half-maximal effective concentration (EC50). The secreted amount of each cytokine in bacterial culture supernatant was measured by ELISA (DuoSet, R&D Systems) as described above (mean of n = 4 replicates: IL-2, 23.0; IL-7, 0.76; IL-15, 1.42; IL-21, 0.86 ng/ml). The EC50 was the value reported for the corresponding recombinant cytokine in a cell proliferation assay (R&D Systems: IL-2, 0.1 to 0.4; IL-7, 0.15 to 0.3; IL-15, 2 to 15; IL-21, 3 to 30 ng/ml), taken as the midpoint of the stated range. Because BLL-IL15 delivers IL-15 genetically fused to the IL-15Rα sushi domain (an RLI-type superagonist), the IL-15 EC50 was divided by 500 to reflect the ∼500-fold greater potency of this fusion over unmodified IL-15 reported for the equivalent construct^32^.

Tumour growth rate was determined for each animal as the mean of the tumour volume changes between consecutive measurement time points divided by the elapsed time (ΔV/Δt) across the treatment course (days 0 to 18). Group mean tumour growth rate was then compared with each payload’s secreted amount relative to its EC50 to assess whether in vitro payload activity was associated with in vivo tumour control.

### Statistical analysis

Statistical tests were performed in GraphPad Prism (version 11.0.2). Group comparisons were made by one-way or two-way ANOVA with correction for multiple comparisons for normally distributed data, or by the Mann–Whitney test as indicated. Survival was analysed by the log-rank test. The details of the statistical tests, including n numbers and P values, are indicated in the respective figure legends. *P* < 0.05 was considered statistically significant.

### Data availability

All data supporting the findings of this study are available within the paper and its Supplementary Information. Bacterial strains and plasmids generated in this study are available from the corresponding author under a standard material transfer agreement.

### Code availability

No custom code was generated in this study. All statistical analyses were performed in GraphPad Prism (version 11.0.2).

## Acknowledgements

We thank members of the Danino laboratory for helpful discussions and feedback on the manuscript. We thank the Biomarkers Core Laboratory (BCL) at the Irving Institute for Clinical and Translational Research, Columbia University, for performing the Luminex multiplex cytokine assays. This publication was supported by the National Center for Advancing Translational Sciences, National Institutes of Health, through Grant Number UL1TR001873. This work was funded by National Institutes of Health R01EB034302 (T.D.), National Science Foundation Graduate Research Fellowship Program award DGE-2036197 (S.W.), Herbert Irving Comprehensive Cancer Center for the Cancer Center Training Grant (A.A.), Pershing Square Foundation (PSF) PSSCRA CU20-0730 (T.D.), Cancer Research Institute (CRI) CRI 3446 (T.D.), Tina’s Wish Foundation TBF CU23-2372 (T.D.), and the Charlie Teo Research Rebel Grant CHTF CU24-1208 (S.W., A.A., T.D.).

## Author contributions

J.I. and T.D. conceived and designed the study. J.I., M.K., H.K., X.H., Y.J., and H.J. designed and constructed the bacterial strains, gene circuits, and payload constructs. J.I., M.K., H.K., and X.H. performed in vitro experiments. J.I., M.K., H.K., X.H., and C.C. performed in vivo tumour studies. S.W. and A.A. performed the intracranial glioblastoma studies, supervised by K.L., J.B., P.C., and T.D. M.A. and K.C. performed the disseminated ovarian cancer studies, supervised by R.D. and T.D. J.I. and M.K. analysed the data. J.I. and T.D. wrote the manuscript with input from all authors.

## Additional information

Supplementary Information is available for this paper.

Correspondence and requests for materials should be addressed to Tal Danino.

## Extended Data

**Extended Data Fig. 1.**
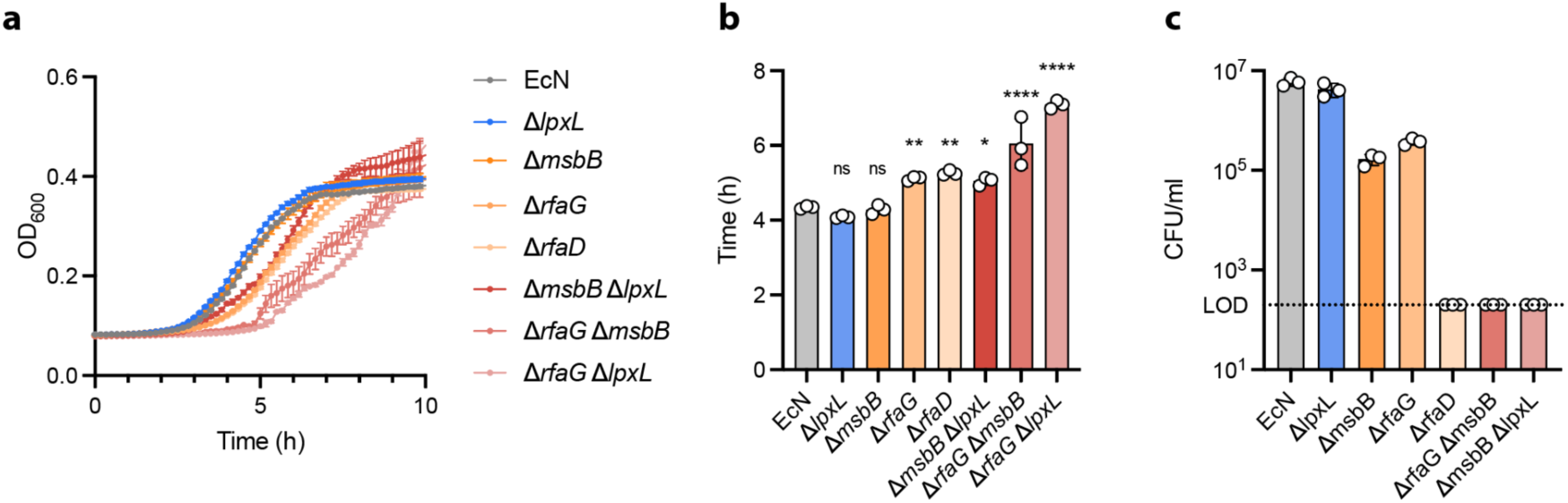
| LPS biosynthetic gene deletions differentially impact EcN growth kinetics and blood survival in vitro. **a**, Growth curves of EcN and LPS deletion mutants measured by optical density at 600 nm (OD_600_) over 10 hours, starting at OD_600_ = 0.01. Data represent mean ± s.e.m. (n = 3 biological replicates). **b**, Time to reach OD_600_ = 0.2 for each strain, interpolated from sigmoidal curve fits in **a**; bars show mean ± s.d. (n = 3 biological replicates). Δ*lpxL* and Δ*msbB* were not significantly different from EcN, whereas Δ*rfaG*, Δ*rfaD*, and all double-deletion strains showed significantly delayed growth. Statistical comparisons versus EcN by one-way ANOVA with Dunnett’s multiple comparisons test (ns, not significant; \**P* < 0.05; \*\**P* < 0.01; \*\*\*\**P* < 0.0001). **c**, Survival of EcN and LPS deletion mutants in human whole blood at 30 minutes post-inoculation, quantified as CFU/ml. Double-deletion strains were cleared to below the limit of detection (LOD, dotted line), whereas single-deletion strains showed variable survival. Data represent mean ± s.d. (n = 3 biological replicates).

**Extended Data Fig. 2.**
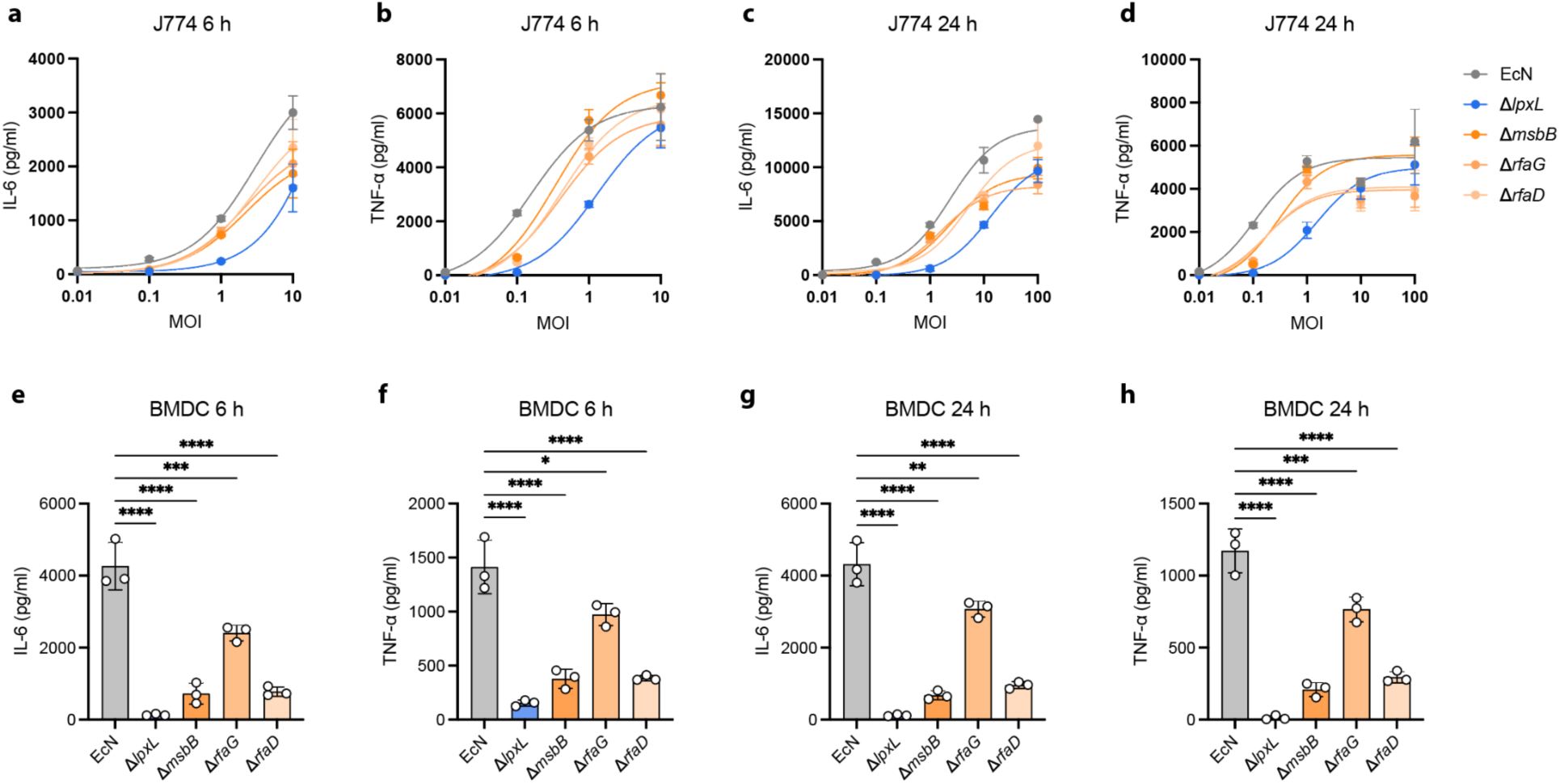
| LPS mutant strains elicit reduced pro-inflammatory cytokine responses in macrophages and bone marrow-derived dendritic cells. **a, b**, IL-6 (a) and TNF-α (b) secretion by J774 murine macrophages stimulated with wild-type EcN or LPS mutant strains across MOIs of 0.01 to 10 for 6 hours. c, d, IL-6 (c) and TNF-α (d) across MOIs of 0.01 to 100 for 24 hours. All mutants showed dose-dependent cytokine induction but remained below wild-type EcN levels, with Δ*lpxL* consistently eliciting the lowest responses. e, f, IL-6 (e) and TNF-α (f) secretion by murine bone marrow-derived dendritic cells (BMDCs) stimulated with wild-type EcN or LPS mutant strains at MOI 0.1 for 6 hours. g, h, IL-6 (g) and TNF-α (h) after 24 hours. All LPS mutants triggered significantly reduced cytokine production compared with wild-type EcN at both time points, with Δ*lpxL* inducing the lowest levels of both cytokines, consistent with the attenuation observed in J774 murine macrophages (Fig. 1e, f). Data represent mean ± s.d. of n = 3 biological replicates. Statistical significance in e–h determined by one-way ANOVA with Tukey’s multiple comparisons test; \**P* < 0.05, \*\**P* < 0.01, \*\*\**P* < 0.001, \*\*\*\**P* < 0.0001.

**Extended Data Fig. 3.**
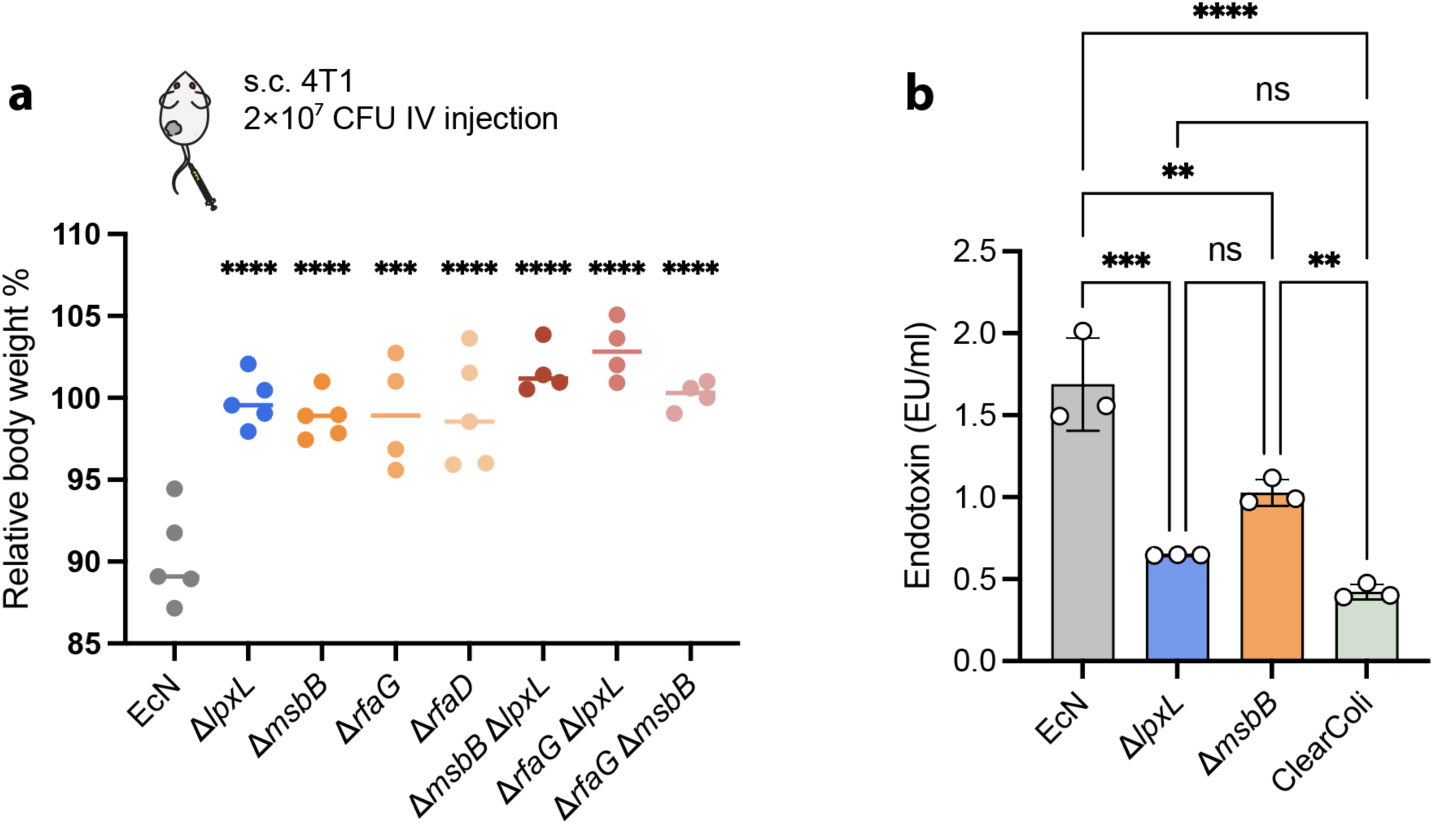
| LPS deletion mutants exhibit improved in vivo tolerability and reduced endotoxin activity compared with EcN. **a**, Relative body weight of 4T1 tumour-bearing BALB/c mice at day 3 following intravenous administration of 2×10^7^ CFU, normalized to day 0. Data represent individual mice (n = 4–5 per group), with the median shown. Asterisks indicate comparisons versus EcN. **b**, Endotoxin activity of whole bacteria (10^3^ CFU per well) quantified by LAL assay. ClearColi served as a low-endotoxin reference. Comparisons are indicated by the connecting lines. Data represent mean ± s.d. (n = 3 biological replicates). Statistical significance by one-way ANOVA with Tukey’s multiple comparisons test (ns, not significant; \*\**P* < 0.01; \*\*\**P* < 0.001; \*\*\*\**P* < 0.0001).

**Extended Data Fig. 4.**
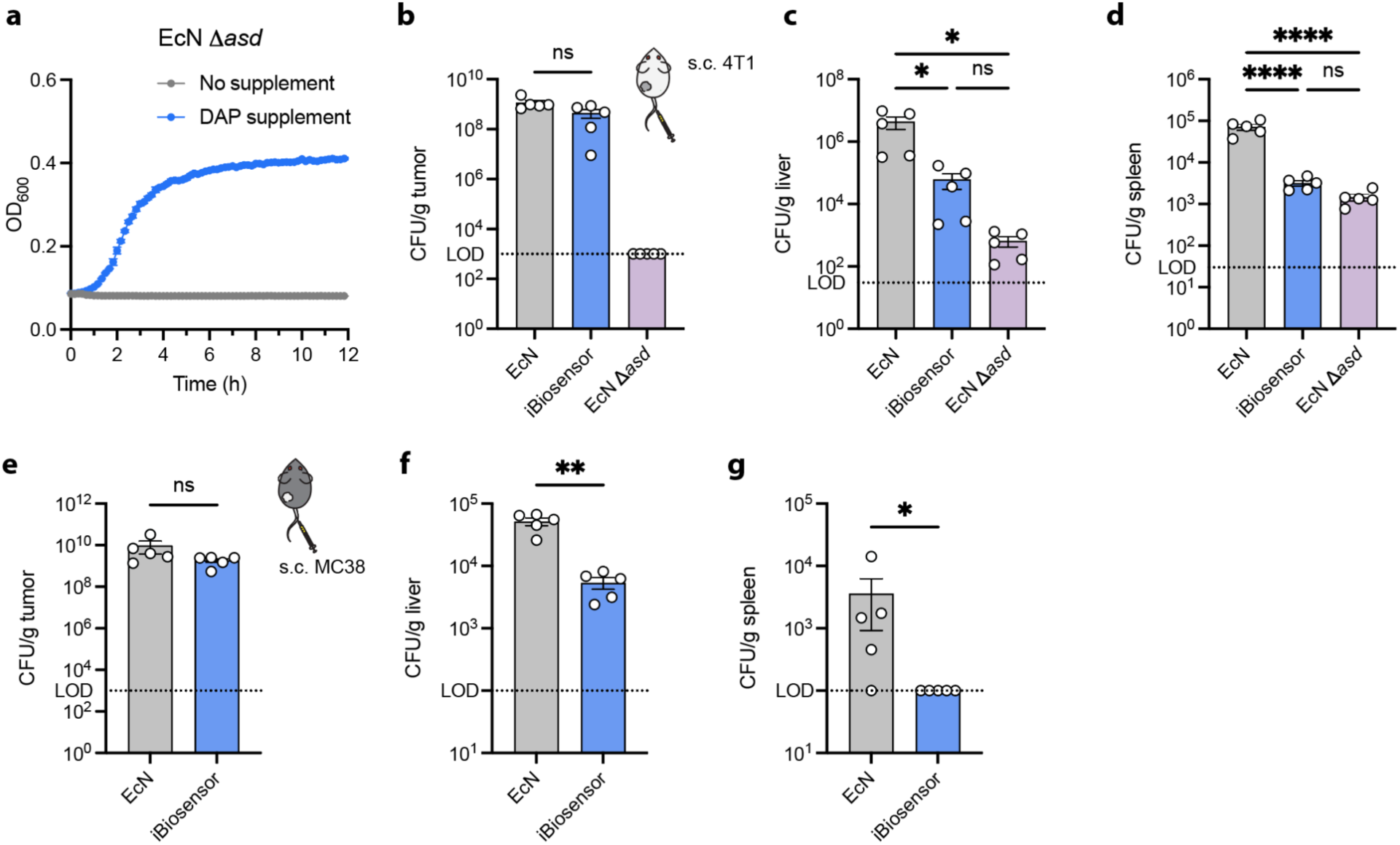
| Auxotrophic EcN Δasd requires DAP supplementation, and biodistribution of EcN, iBiosensor, and EcN Δasd strains. **a**, Growth curves of EcN Δ*asd* in the presence or absence of DAP, measured by OD_600_ over 12 hours. Δ*asd* fails to grow without DAP supplementation, confirming functional auxotrophy. Data represent mean ± s.e.m. (n = 3 biological replicates). b–d, Bacterial colonization in tumour (b), liver (c), and spleen (d) (CFU/g tissue) of 4T1-bearing BALB/c mice at day 1 following i.v. injection of 4×10^7^ CFU. EcN Δ*asd* serves as an auxotrophic control lacking the ability to grow. e–g, Bacterial colonization in tumour (e), liver (f), and spleen (g) (CFU/g tissue) of MC38-bearing C57BL/6 mice at day 2 following i.v. injection of 1×10^7^ CFU. The dotted line indicates the limit of detection (LOD). Data are mean ± s.e.m. (n = 5 per group). Statistical significance by one-way ANOVA with Tukey’s multiple comparisons test (b–d) or Mann–Whitney test (e–g); \**P* < 0.05, \*\**P* < 0.01, \*\*\*\**P* < 0.0001; ns, not significant.

**Extended Data Fig. 5.**
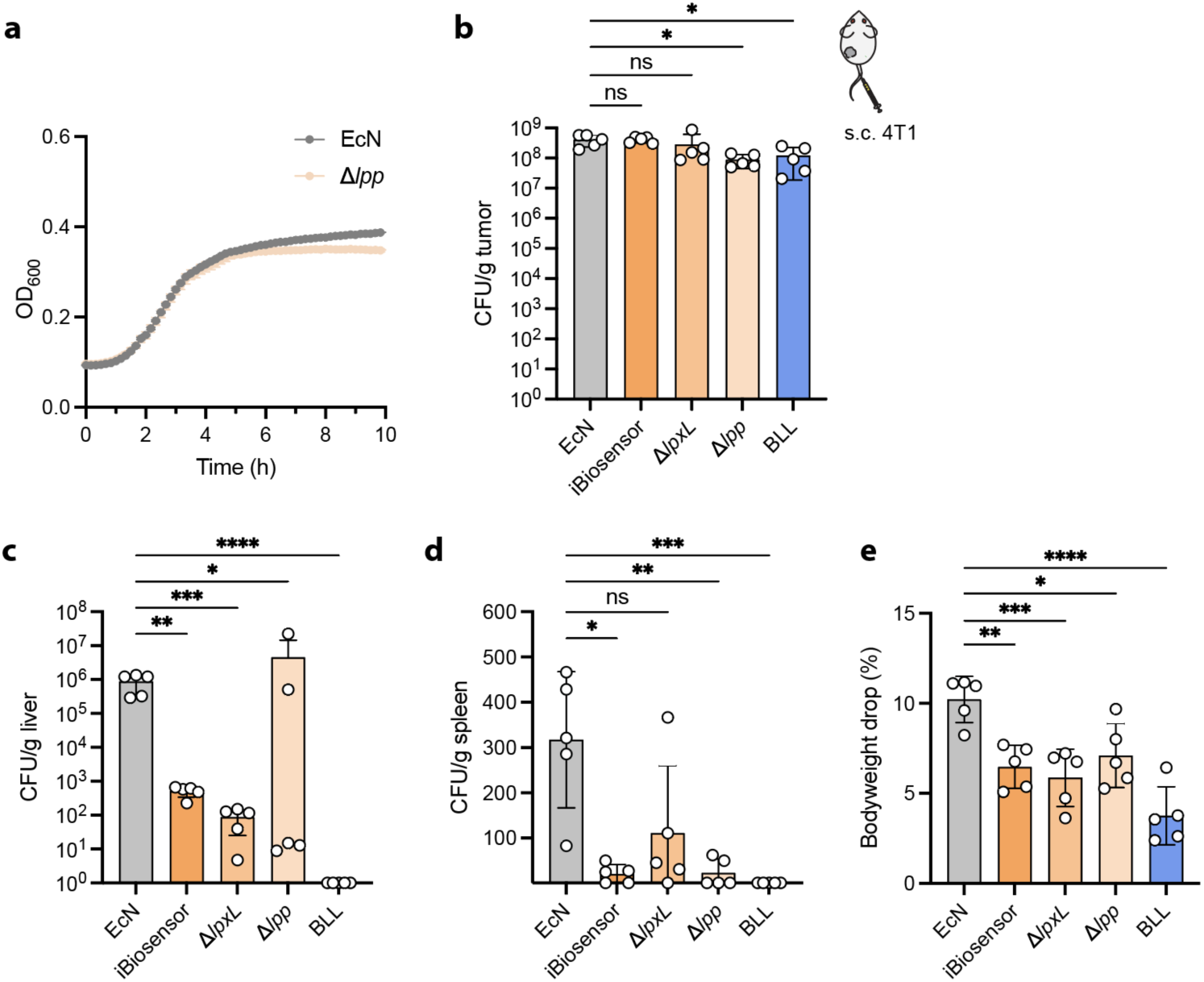
| Growth kinetics of EcN Δlpp, and biodistribution and tolerability of individual EcN modifications and BLL in BALB/c mice. **a**, Growth curves of wild-type EcN and EcN Δ*lpp* in LB medium over 10 hours. EcN Δ*lpp* exhibits growth kinetics comparable to wild-type EcN. Data are mean ± s.e.m. (n = 3 biological replicates). b, Tumour, c, liver, d, spleen colonization, and e, day 1 body weight drop after intravenous injection of 2×10^7^ CFU of EcN, iBiosensor, Δ*lpxL*, Δ*lpp*, or BLL in 4T1 tumour-bearing BALB/c mice (n = 5 mice per group). All strains colonize tumours at comparable levels, whereas BLL shows markedly reduced liver and spleen colonization and minimal body weight loss relative to wild-type EcN. Colonization data (b–d) are geometric mean ± geometric s.d.; body weight data (e) are mean ± s.d.. Statistical significance by lognormal one-way ANOVA with Dunnett’s multiple comparisons test (b–d) or ordinary one-way ANOVA with Dunnett’s multiple comparisons test (e); \**P* < 0.05, \*\**P* < 0.01, \*\*\**P* < 0.001, \*\*\*\**P* < 0.0001; ns, not significant.

**Extended Data Fig. 6.**
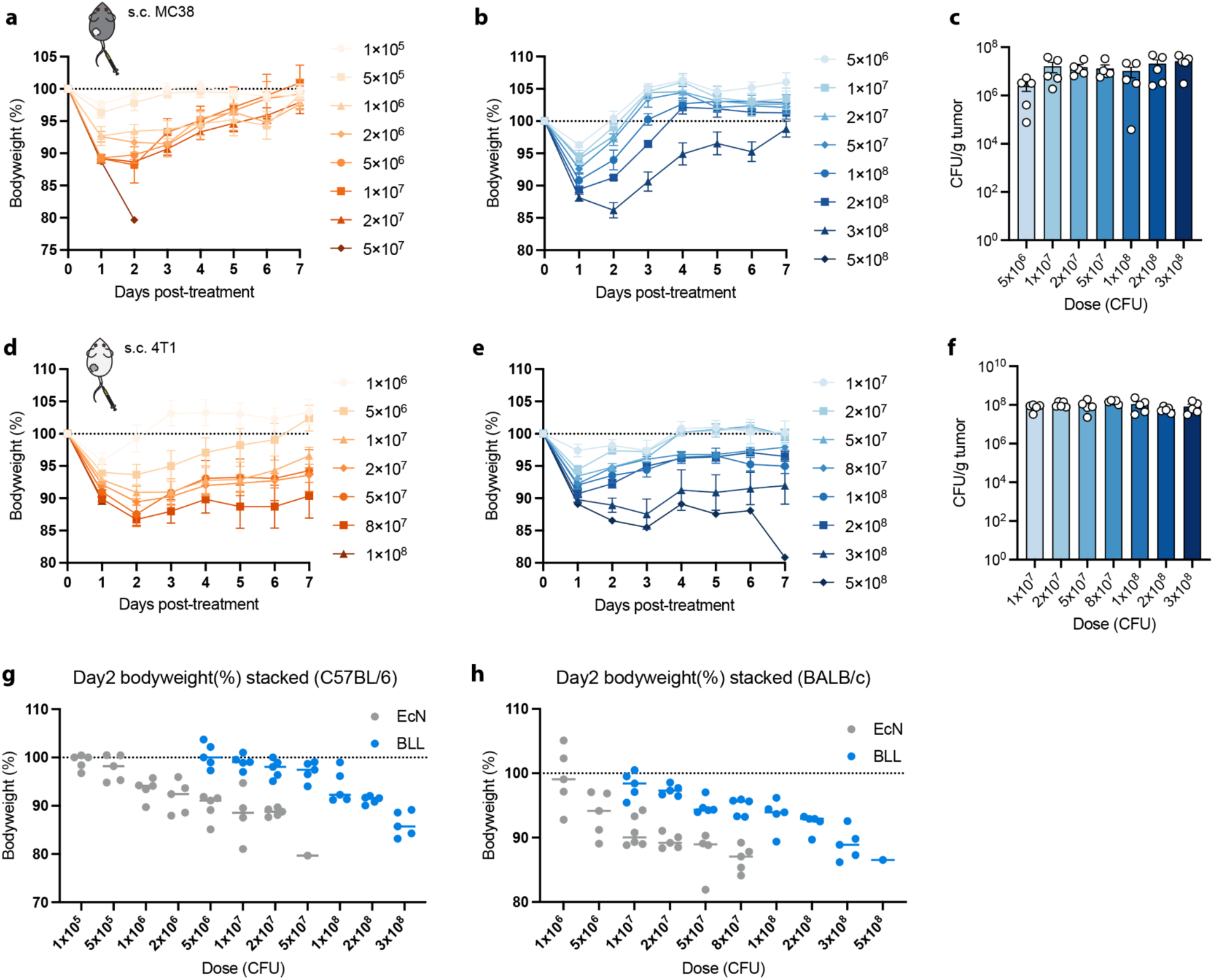
| Dose-escalation tolerability and tumour colonization across two syngeneic mouse backgrounds. **a**–**c**, MC38 tumour-bearing C57BL/6 mice. **a**, Body weight over 7 days after intravenous EcN across escalating doses of 1×10^5^ to 5×10^7^ CFU (orange). **b**, Body weight over 7 days after intravenous BLL across escalating doses of 5×10^6^ to 5×10^8^ CFU (blue). **c**, Intratumoural BLL burden at day 7 across doses, showing dose-independent colonization. **d**–**f**, 4T1 tumour-bearing BALB/c mice. **d**, Body weight after intravenous EcN across 1×10^6^ to 1×10^8^ CFU (orange). **e**, Body weight after intravenous BLL across 1×10^7^ to 5×10^8^ CFU (blue). **f**, Intratumoural BLL burden at day 7 across doses. **g**, **h**, Day 2 body weight of EcN (grey) and BLL (blue) overlaid at matched doses, illustrating the expanded therapeutic window of BLL in the C57BL/6 (**g**) and BALB/c (**h**) backgrounds. Body weight data in **a**, **b**, **d**, and **e** are mean ± s.e.m.. In **c** and **f**, bars represent mean ± s.e.m. and symbols represent individual mice. In **g** and **h**, horizontal bars represent the median and symbols represent individual mice.

**Extended Data Fig. 7.**
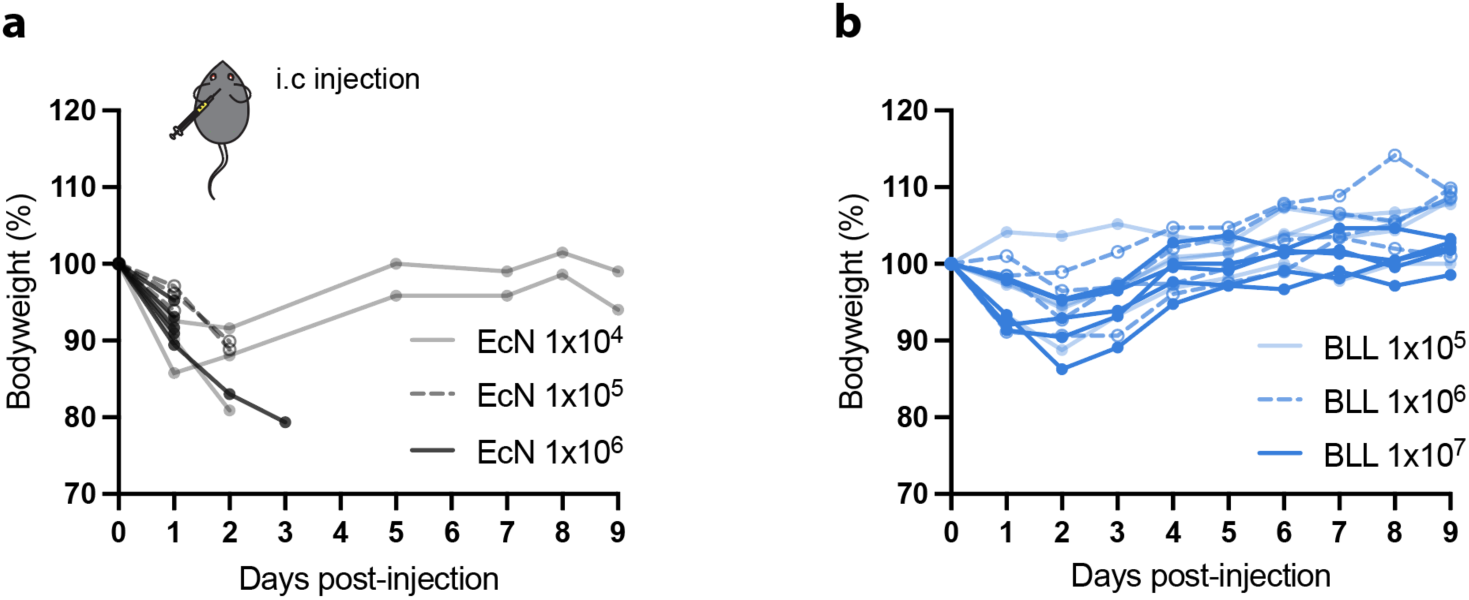
| BLL is tolerated at higher intracranial doses than EcN. **a**, **b**, Relative body weight of C57BL/6 mice following intracranial injection of EcN (**a**) or BLL (**b**) at escalating doses. EcN caused acute weight loss with early mortality at all doses tested, whereas BLL was tolerated across the dose range with recovery to baseline by day 9. **a**, EcN: n = 3 mice for 1×10^4^ and 1×10^5^, n = 6 for 1×10^6^. **b**, BLL: n = 4 mice for 1×10^5^ and 1×10^6^, n = 5 for 1×10^7^.

**Extended Data Fig. 8.**
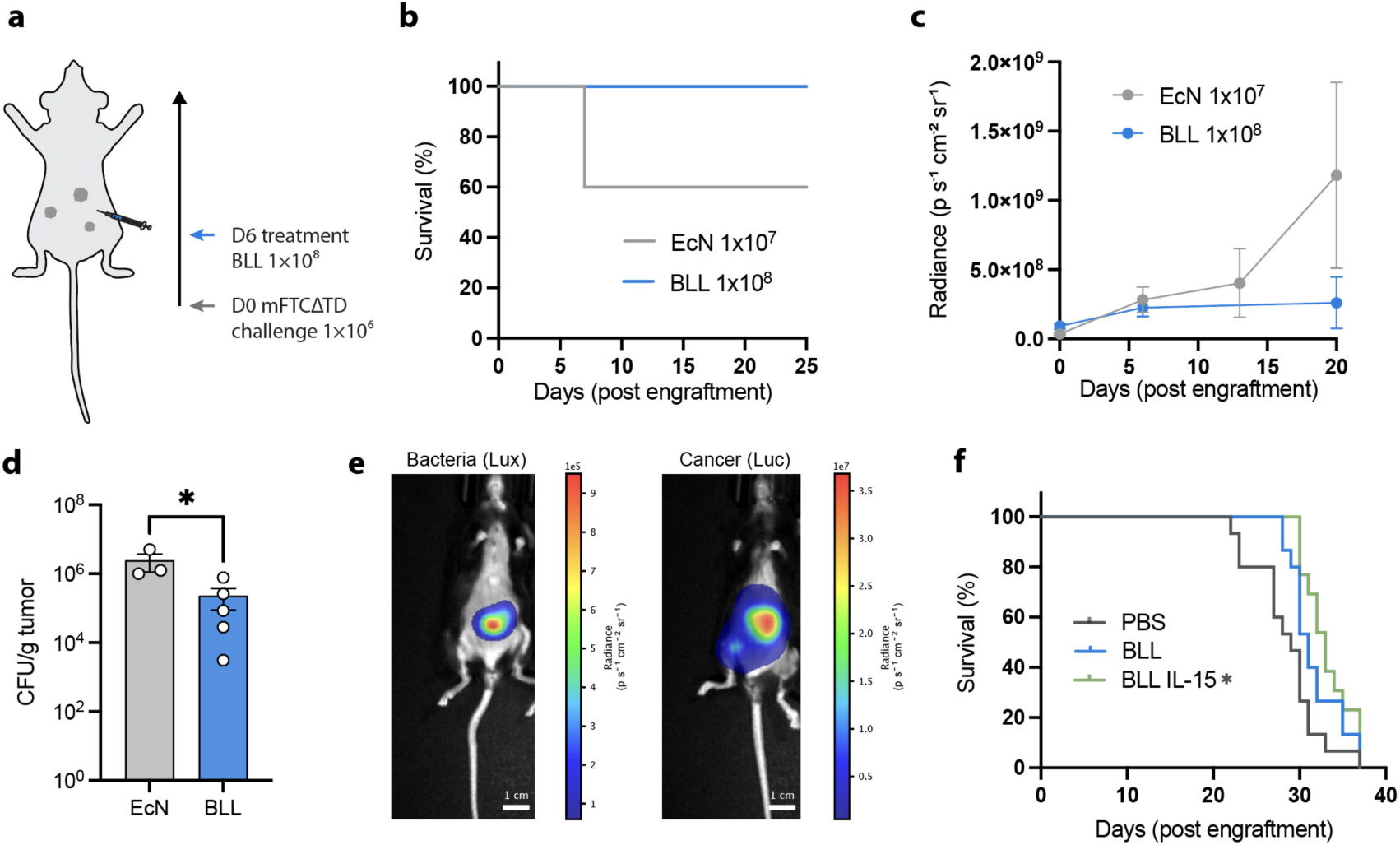
| BLL enables safe intraperitoneal treatment and IL-15-mediated survival benefit in a disseminated ovarian cancer model. **a**, Experimental schematic for b–e. C57BL/6 mice were challenged intraperitoneally (i.p.) with 1×10^6^ mFTC-ΔTD-luc cells on day 0, followed by i.p. bacterial treatment on day 6 with EcN (1×10^7^ CFU) or BLL (1×10^8^ CFU); n = 5 mice per group. b, Kaplan–Meier survival curves. EcN-treated mice (1×10^7^ CFU) exhibited treatment-related mortality (2 of 5 mice), whereas BLL-treated mice (1×10^8^ CFU, 10-fold higher dose) maintained 100% survival through day 25 (n = 5 per group). c, Tumour burden over time measured by bioluminescence imaging (BLI) of mFTC-ΔTD-luc cells (radiance, p s^−1^ cm^−2^ sr^−1^). BLL-treated mice showed reduced tumour progression compared with EcN-treated mice. d, Tumour colonization (CFU/g tumour) at the study endpoint (day 29). e, Representative bioluminescence images at day 20. Left, bacterial luminescence (BLL expressing the lux operon). Right, tumour luminescence (mFTC-ΔTD-luc). Scale bar, 1 cm. f, Kaplan–Meier survival curves from an independent experiment in the same mFTC-ΔTD model, in which mice received intravenous injection of PBS, BLL, or BLL-IL15 (1×10^8^ CFU) on day 1 post-engraftment (n = 15, 15, and 13, respectively). BLL-IL15 significantly extended survival relative to PBS. Data are mean ± s.e.m. (c, d). Statistical significance by Mann–Whitney test (d) or log-rank (Mantel-Cox) test with Šídák correction for multiple comparisons (f); \**P* < 0.05.

**Extended Data Fig. 9.**
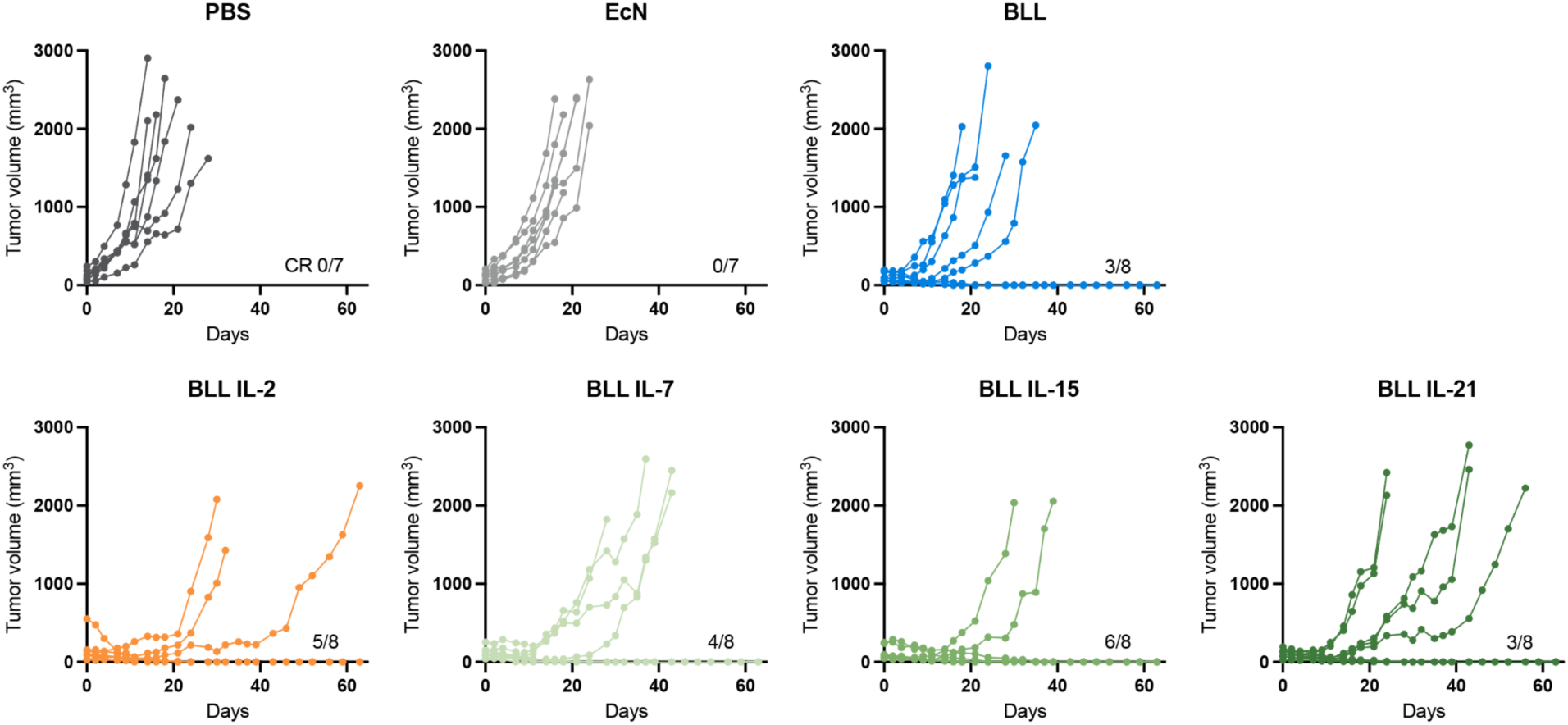
| Individual tumour growth trajectories in the MC38 model. Individual tumour volumes over time for PBS (0 of 7 CR), EcN (0 of 7 CR), BLL (3 of 8 CR), BLL-IL2 (5 of 8 CR), BLL-IL7 (4 of 8 CR), BLL-IL15 (6 of 8 CR), and BLL-IL21 (3 of 8 CR). CR, complete response, defined as tumour volume = 0 mm^3^ sustained through day 60.

**Extended Data Fig. 10.**
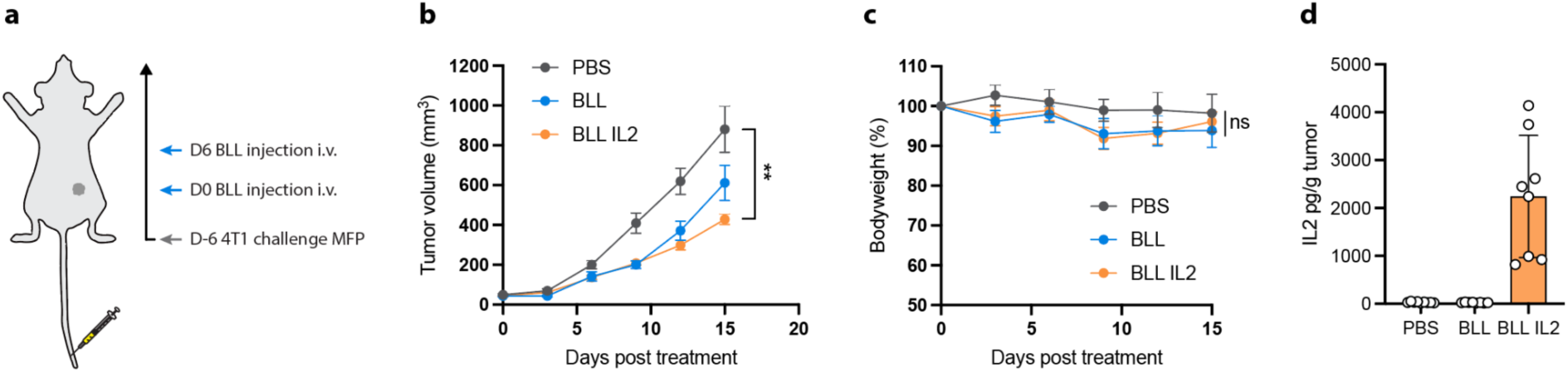
| BLL-IL2 reduces tumour growth in the 4T1 mammary carcinoma model. **a**, Experimental schematic. BALB/c mice bearing orthotopic 4T1 mammary fat pad tumours, challenged on day −6, received intravenous treatment on days 0 and 6. **b**, Mean tumour volume over time following i.v. treatment with PBS, BLL, or BLL-IL2 at the maximum tolerated dose in BALB/c mice bearing orthotopic 4T1 mammary fat pad tumours (n = 10, 9, 10). **c**, Relative body weight over the treatment period. **d**, Intratumoural IL-2 concentration (pg/g tumour) at endpoint. Data are mean ± s.e.m. (**b**, **c**) or mean ± s.d. (**d**). Statistical significance by two-way ANOVA with Tukey’s multiple comparisons test at the final timepoint (**b**, **c**); \*\**P* < 0.01; ns, not significant.

